# Unveiling the native architecture of adult cardiac tissue using the 3D-NaissI method

**DOI:** 10.1101/2024.11.26.625417

**Authors:** Nicolas Pataluch, Céline Guilbeau-Frugier, Véronique Pons, Amandine Wahart, Clément Karsenty, Jean-Michel Sénard, Céline Galés

## Abstract

Accurately imaging adult cardiac tissue in its native state is essential for regenerative medicine and understanding heart disease. Current fluorescence methods encounter challenges with tissue fixation. Here, we introduce the 3D-NaissI (3D-Native Tissue Imaging) method, enabling rapid, cost-effective imaging of fresh cardiac tissue samples in their closest native state, that we also extended to other tissues.

We validated 3D-NaissI’s efficacy in preserving cardiac tissue integrity using small biopsies under hypothermic conditions in phosphate-buffered saline, offering unparalleled resolution in confocal microscopy for imaging fluorescent-small molecules/-antibodies. Compared to conventional histology, 3D-NaissI preserves cardiac tissue architecture and native protein epitopes, facilitating the use of a wide range of commercial antibodies without unmasking strategies. We successfully identified specific cardiac protein expression patterns in cardiomyocytes (CMs) from rodents and humans, including for the first time ACE2 localization in the lateral membrane/T-Tubules and SGTL2 in the sarcoplasmic reticulum. These findings shed light on COVID-19-related cardiac complications and suggest novel explanations for iSGLT2 therapeutic benefits in HFpEF patients. Additionally, we challenge the notion of "connexin-43 lateralization” in heart pathology, suggesting it may be an artifact of cardiac fixation, as 3D-NaissI clearly revealed native connexin-43 expression at the lateral membrane of healthy CMs. We also discovered previously undocumented periodic ring-like 3D structures formed by pericytes covering CMs’ lateral surfaces. These structures, positive for laminin-2, delineate a specific spatial architecture of laminin-2 receptors at the CM surface, highlighting the pivotal role of pericytes in CM function. Lastly, 3D-NaissI facilitates mapping native human protein expression in fresh cardiac autopsies, providing insights into both pathological and non-pathological contexts.

Hence, 3D-NaissI offers unparalleled insights into native cardiac tissue biology and promises to advance our understanding of physiology and pathophysiology, surpassing standard histology in resolution and accuracy.

## Introduction

The quest for successfully regenerating adult cardiac tissue, a paramount objective in treating end-stage heart failure, has primarily centered around cell-dependent strategies. Despite early optimism, efforts such as stem cell engraftment and enhancing resident cardiomyocyte (CM) proliferation have fallen short, often leading to arrhythmia and insufficient functional recovery[1–3]. In contrast, 3D cardiac tissue engineering, considering the complex interplay between diverse cell populations and the extracellular matrix, emerges as a promising alternative. However, its potential hinges on clinical validation, necessitating further research. Paradoxically, while extensive studies focus on regenerating pathological cardiac tissue, our understanding of the 3D organization of native cardiac tissue remains a mystery - a crucial gap for engineering functional cardiac tissue faithful to native conditions.

Cardiology’s insights into the 3D architecture of the mammalian heart largely rely on noninvasive macroscopic imaging like X-rays and echocardiography, valuable for overall structure but lacking at the microscopic level. Recent advances in tissue clearing techniques[4–7], though significant, raise concerns about preserving native tissue architecture due to chemical fixation risks. Conventional fixatives like paraformaldehyde (PFA) and glutaraldehyde (GA) pose risks of altering tissue biochemistry, inducing artifacts, and causing epitope masking, particularly challenging in solid cohesive tissues. Crosslinking during fixation (protein-protein, DNA-DNA, DNA-RNA, or DNA-protein crosslinking) can lead to various artifacts, impacting not only tissue-clearing but also histopathology methods, considered the gold standard for clinical diagnosis. Mutational signatures associated with formalin fixation on patient samples[8] and alterations in tissue biochemistry revealed by Raman spectroscopy (https://doi.org/10.1002/jrs.4223) highlight concerns in conventional histopathology examination. Moreover, epitope masking caused by fixatives, a well-known challenge in histopathology[9], can hinder protein recognition in immunodetection methods, especially in solid cohesive tissues. Fixation artifacts are also observed in electron microscopy, necessitating reliable artifact-free methods for tissue examination. Fixation’s influence on protein localization, demonstrated by recent studies[10], emphasizes the need for live-cell imaging techniques to closely resemble the native state[11]. Despite their known limitations, fixation methods, developed to capture a snapshot of cells or tissues, extensively persist in cell biology without reassessing the native biological sample, lacking means to image tissues in their closest native state.

Addressing these limitations, we introduce 3D-NaissI (**Na**tive Ti**ss**ue **I**maging), a novel method for fresh-tissue immunolabeling applicable to small biopsies, allowing fluorescent labeling without fixation or permeabilization. Beyond cardiac tissue, we also validated its utility to brain, skin, and gut biopsies. Rapid, cost-effective, and autofluorescence-free, 3D-NaissI identifies previously overlooked 3D structures within cardiac tissue. It enabled high-resolution confocal imaging of traditionally challenging cardiac proteins, surpassing formalin-fixed processing. When compared to standard histological methods, 3D-NaissI exposed interpretation errors, challenging established ideas in cardio-pathology, such as the concept of "Cx43 lateralization. Finally, applied successfully in cardiac autopsies, 3D-NaissI facilitates mapping native protein expression in the human heart with minimal artifacts aside from the patient’s clinical data. This innovative approach promises to fill the existing gap by providing a means to image tissues in their closest native state, significantly advancing our understanding and applications in the field of tissue imaging.

## Results

### Fresh native cardiac tissue biopsies: preparation, staining and imaging

The current gold standard for *ex vivo* cardiac tissue culture relies on living myocardial slices (LMS), optimized for functional studies to maintain adult cardiac tissue for extended periods[12]. However, this conditioning is geared towards functional assessments, such as contraction and electrical conduction, rather than imaging. Typically, imaging is performed using PFA-fixed LMS cryosections[13, 14].

In this study, our objective was to develop a technique that optimally preserves cardiac tissue integrity for short-term imaging in a state closely resembling its native condition, eliminating the need for functional preservation. Hence, we developed a protocol referred to as the 3D-NaissI (Native tissue Imaging) method (**Fig. 1a**). Cardiac tissue sampling from mice followed a previously described approach for preserving the subcellular architecture of cardiomyocytes (CMs) in electron microscopy imaging[15]. Smoothly cut 1-mm^3^ fresh biopsies were rapidly collected from the left ventricular myocardium of mice under hypothermic conditions at 4°C (**Supplementary Fig. 1**) and maintained in phosphate-buffered saline (PBS) at 4°C until imaging (**Fig. 1a**). The small biopsy size is crucial for subsequent tissue penetration of fluorescent probes, although it may lead to section artifacts (tissue tearing), necessitating the preparation of multiple biopsies from each sample.

**Figure 1.**
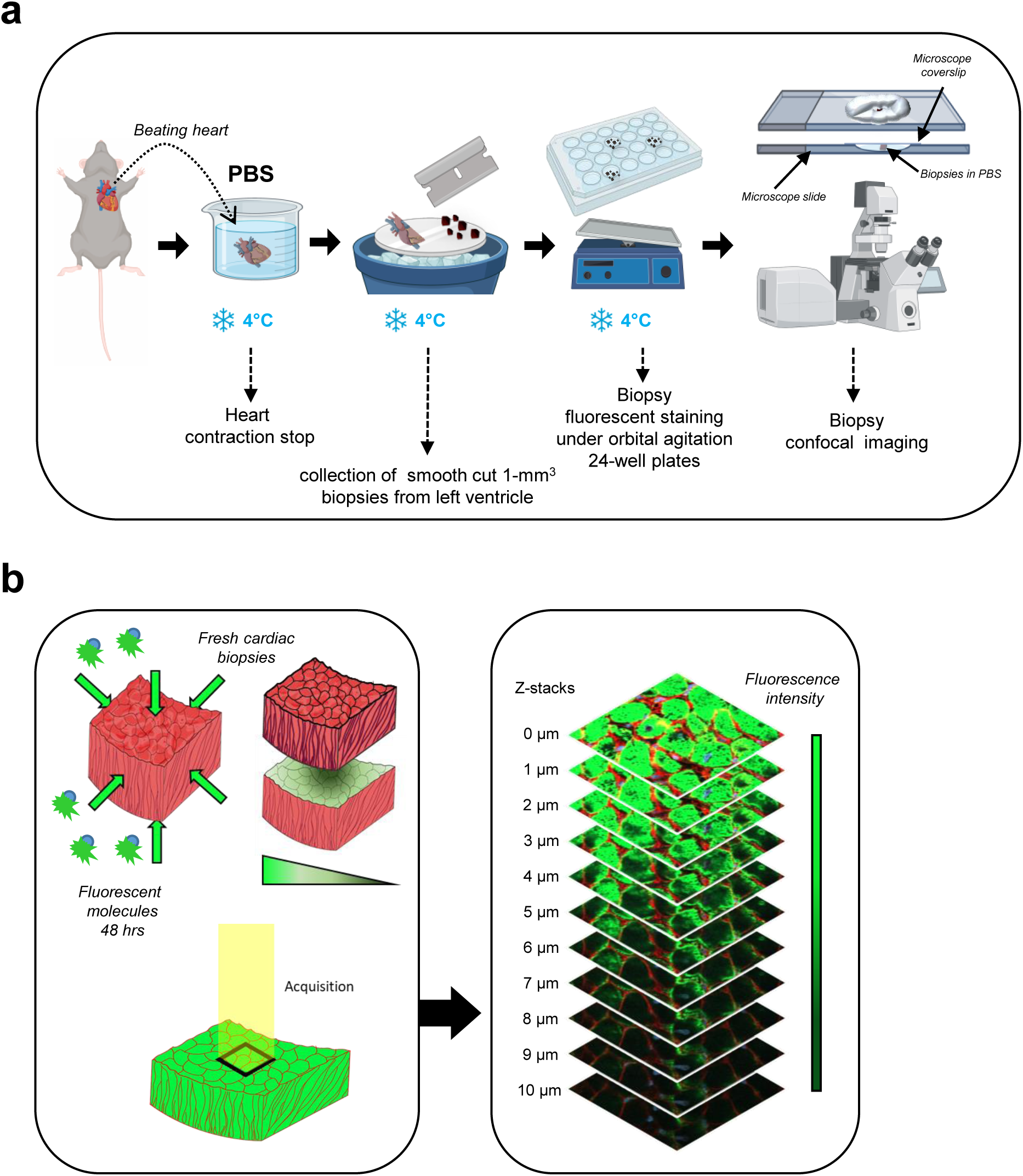
Schematic workflow of 3D-NaissI method. **a** Procedure from beating heart retrieval to biopsy preparation for confocal imaging. **b** Confocal imaging of fluorescent staining in fresh cardiac biopsies and z-stack acquisitions.

We first conducted a thorough qualitative and quantitative evaluation of staining solutions on fresh biopsies at different temperatures. Direct fluorescent staining using small fluorescent molecules was prioritized for their enhanced tissue penetration. Fresh cardiac biopsies were incubated for 48 hours in the dark with agitation in 24-well plates, using PBS, Hanks’ balanced salt solution (HBSS), or a cardioplegic solution at 4°C, 22°C (room temperature), or 37°C. Stained biopsies were mounted and immobilized for confocal imaging. To prevent tissue edge damages, the top layers of the biopsies were excluded from image acquisition (See Methods), and 1 µm stacks were acquired until staining loss occurred (**Fig. 1b**). Related Z-stacks were compared among the different samples. Fluorescent staining may exhibit heterogeneity due to the small size and non-oriented nature of the biopsies (**Supplementary Fig. 2**), necessitating multiple samples per condition for a comprehensive view of the tissue. The cytoarchitecture of the CMs, reflected by both actin (SPY^TM^-actin cell permeable-live cell fluorescent probe) and Wheat-Germ-Agglutinin (WGA)-surface stainings, was well-preserved at 4°C, regardless of the buffer used (**Fig. 2a**). At 22°C, tissue disorganization had already begun, hindering fluorescent staining. At 37°C, extensive damage occurred in all buffers, rendering the tissue unsuitable for further use. These findings were further supported by a quantitative assessment of the CM inter-lateral space area (**Fig.2b**) (See Methods), which typically expands in pathophysiology due to cardiac tissue disorganization and resulting in a loss of tissue cohesion[16]. Due to the variability in fluorescent staining among different buffers, only a few PBS quantifications were performed at 22°C. The results showed a significantly higher space at 22°C compared to 4°C in PBS (**Fig. 2b**, **left panel**), indicating loss of tissue cohesion. Conversely, the inter-cellular space at 4°C remained well-maintained across all preservation buffers, albeit with a slight increase observed in HBSS and the cardioplegic condition (**Fig. 2b**, **right panel**). Larger cardiac biopsies could also be processed in PBS at 4°C for confocal fluorescence imaging (**Supplementary Fig. 3**).

**Figure 2.**
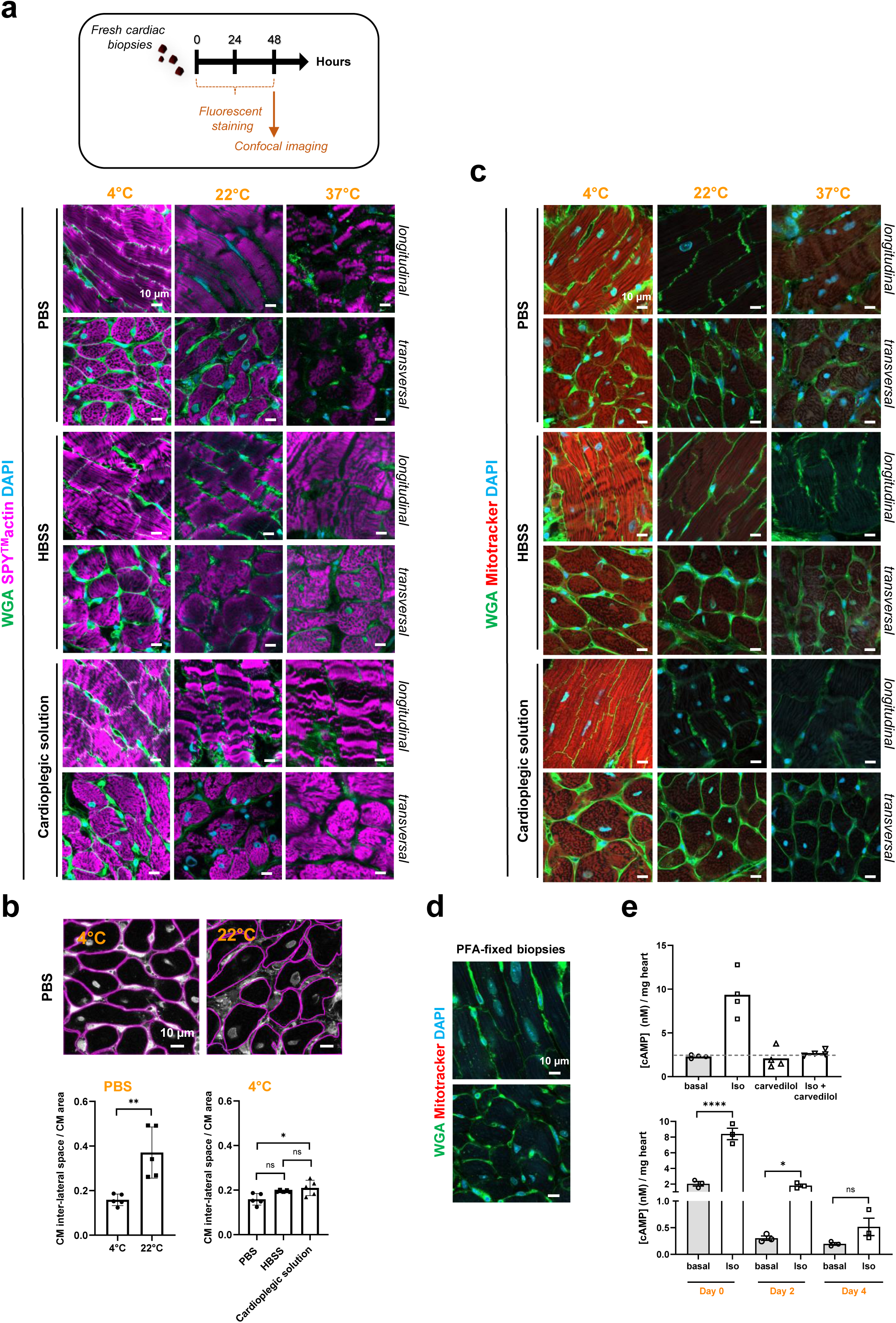
Optimizing experimental conditions to preserve stability, integrity, and viability of fresh cardiac tissue biopsies using the 3D-NaissI method. **a** Stability-integrity of fresh cardiac biopsies from mice qualitatively evaluated by confocal imaging following 48 hours of fluorescent staining with cell surface probe (wheat germ agglutinin-WGA), cytoskeletal actin (SPY^TM^-actin) and nuclei (DAPI) at 4, 22 or 37 °C and in PBS, HBSS or a cardioplegic solution. Images illustrate cardiomyocyte (CM) cytoarchitecture under the different conditions and are representative of 3-5 independent experiments (5 mice). **b** Stability-integrity of fresh cardiac biopsies assessed by quantification of the inter-CM space (indicative of the cardiac tissue cohesion) in fluorescent-WGA-stained fresh biopsies at 4 or 22°C in PBS, HBSS or a cardioplegic solution. Data are mean ± s.d. *n=5* mice (5-10 biopsies/mouse, 1 image /biopsy), one-way ANOVA, Tukey post-hoc test. **c** Mitochondria viability in fresh cardiac biopsies from mice qualitatively evaluated by confocal imaging following 48 hours of fluorescent staining with cell surface probe (WGA)/Mitochondria membrane potential probe (Mitotracker) and nuclei (DAPI) at 4, 22 or 37 °C and in PBS, HBSS or a cardioplegic solution. Images are representative of 3 independent experiments (3 mice). **d** Negative control for mitotracker probe staining using paraformaldehyde (PFA)-fixed, non-living cardiac biopsies. **e** Viability of fresh cardiac biopsies from mice assessed by measuring β-adrenergic activity of the tissue. ***Upper panel:*** cAMP production quantified in biopsies stimulated or not (basal) with 10 µM isoproterenol (ISO) or 10 µM carvedilol alone or in combination for 30 min at room temperature. Data represent the mean ± s.e.m. of 4 different biopsies from one mouse and are expressed as cAMP concentration (nM)/mg heart. ***Lower panel*:** cAMP production quantified in fresh cardiac biopsies collected immediately (Day 0), or 2 (Day 2), or 4 (Day 4) days after collection and stimulated or not (basal) with 10 µM isoproterenol (ISO) for 30 min at room temperature. Data are mean ± s.e.m., *n=3* mice (3-4 biopsies/mouse), one-way ANOVA, Holm-Sidak’s post-hoc test. (* *P*<0.05; **** *P*<0.0001; ns, not statistically significant).

### Fresh native cardiac tissue biopsies: stability-viability

Fresh cardiac biopsies stored in PBS at 4°C demonstrated sustained tissue stability for at least 10 days, as evidenced by consistent CM inter-lateral space and CM area (**Supplementary Fig. 4a**). Likewise, in a similar setting, sarcomeric α-actinin expression, a constituent of the CM contractile apparatus, remained stable up to 10 days (**Supplementary Fig. 4b**). Optimal preservation of fresh cardiac biopsies at 4°C was further confirmed using the MitoTracker probe (**Fig. 2c**). Positive fluorescent MitoTracker labeling, indicative of viable mitochondria, was observed in biopsies at 4°C in all buffers. In contrast, no labeling was observed at 22°C and 37°C, indicating loss of tissue viability. Notably, mitochondria viability persisted for at least 7 days at 4°C (**Supplementary Fig. 5a**). As expected, MitoTracker fluorescence was absent in PFA-fixed biopsies, confirming loss of mitochondria viability (**Fig.2d**). Similarly, plasma membrane integrity in viable CMs, assessed using the potentiometric fluorescent dye di-8-ANEPPS, remained evident for up to 10 days only in whole biopsy CM (**Supplementary Fig. 5b)**. These results were supported by a quantitative assessment of the β-adrenergic receptor (β-AR)/Adenylyl-cyclase/cAMP signaling pathway (**Fig. 2e**). Stimulation of freshly collected heart biopsies with isoproterenol (nonselective β-agonist) at room temperature for 30 minutes resulted in significant cAMP production, which was completely blocked by co-stimulation with carvedilol (β-antagonist), confirming β-AR dependence (**Fig. 2e**, **upper panel**). This β-AR response, encompassing G protein and adenylyl cyclase enzymatic activity, was maintained in biopsies stored in PBS at 4°C for up to 2 days but was lost after 4 days (**Fig. 2e**, **lower panel**), aligning with short-term preservation of cellular activity under cold exposure. These results were further supported by the rapid decline in GAPDH enzyme expression during the first 2 days in the cardiac tissue samples stored in PBS at 4°C (**Supplementary Fig. 4b**).

In summary, these findings underscore that fresh cardiac biopsies stored and stained at 4°C in PBS exhibit superior preservation of tissue integrity. Notably, this approach effectively maintains both tissue stability and viability, with structural proteins exhibiting greater stability than enzymatic counterparts. These optimized experimental conditions were consistently employed for all subsequent experiments using the 3D-NaissI method.

### 3D-NaissI method: comparison with common histological techniques

To assess the 3D-NaissI method against conventional histological approaches in cardiac tissue fluorescence imaging, we compared WGA and DAPI small-fluorescent molecule staining on fresh tissue (3D-NaissI), PFA-prefixed tissue, and Formalin-Fixed Paraffin-Embedded (FFPE) tissue. Native tissue exhibited minimal autofluorescence across various filters, contrasting with significant autofluorescence in PFA or FFPE tissue (**Supplementary Fig. 6**). These outcomes align with the expected autofluorescent properties resulting from crosslinking with amines and proteins in the presence of PFA or formalin. This discrepancy underscores the 3D-NaissI method’s substantial enhancement of the fluorescent signal-to-noise ratio. Beyond signal quality, using native fresh tissue for cardiac imaging surpasses classical histological methods by better preserving cellular and tissue integrity, as qualitatively shown in **Fig. 3a**. Accordingly, the CM inter-lateral space (**Fig. 3b**, **left panel**) and the CM area (**Fig. 3b**, **right panel**) were significantly smaller in PFA-fixed and FFPE tissues compared to fresh biopsies, indicating compromised tissue cohesion and cellular alterations in the presence of fixatives. These alterations may influence the intricate architecture of CMs, protein expression, and localization, critical in cardiology research. In conclusion, conventional histological techniques considerably shrink cardiac tissue and are not appropriate for precisely studying cellular architecture and spatial organization of proteins *in situ* compared to 3D-NaissI method.

**Figure 3.**
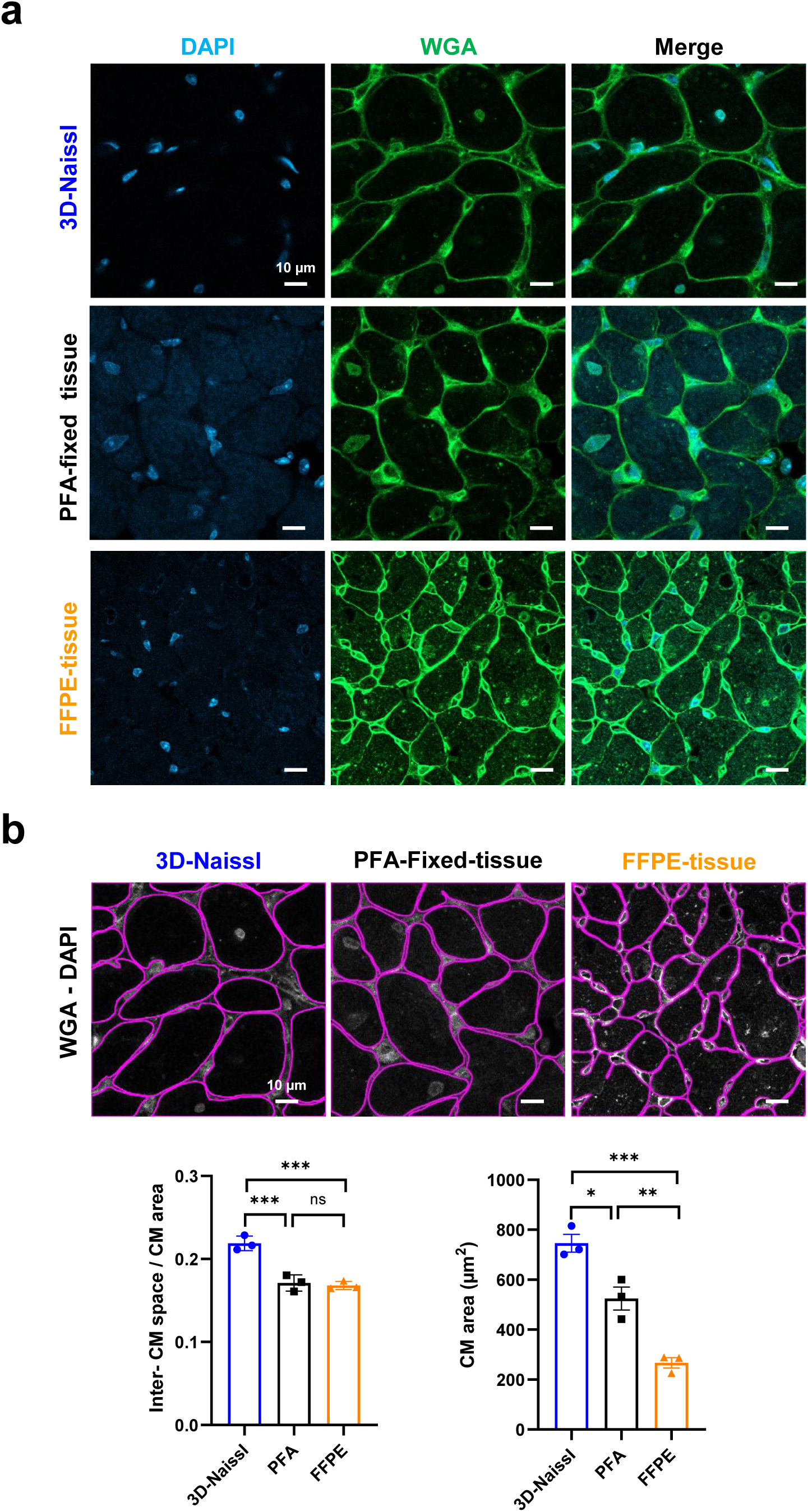
Performance of 3D-NaissI compared to conventional histological techniques. **a** Representative confocal images of fluorescent wheat germ agglutinin (WGA) and DAPI-staining in cardiac tissue (left ventricles) from fresh cardiac biopsies (3D-NaissI method), PFA-fixed-tissue or Formalin-Fixed Paraffin-Embedded (FFPE) tissue. **b** Quantification of inter-CM space (indicative of the cardiac tissue cohesion) and CM area in fluorescent-WGA-stained cardiac tissue using fresh cardiac biopsies (3D-NAissI; PBS, 4°C), PFA-fixed tissue or FFPE-tissue. Data are mean ± s.d., *n*=3 mice (7-10 biopsies/mouse, 1 image /biopsy), one-way ANOVA, Tukey post-hoc test.

### 3D-NaissI method: optimization of fluorescent staining and immunostaining

We optimized fluorescent staining of fresh cardiac tissue at 4°C in PBS using small molecules. A 30-minute labeling allowed visualization of DAPI, Alexa Fluor 488-WGA, and Alexa Fluor 594-mitotracker fluorescence. Extended incubation times (24-48 hours) resulted in an optimal imaging depth of ∼40 µm (**Supplementary Fig. 7**), aligning with confocal microscope capabilities. The signal-to-noise ratio was improved at greater depths by linearly increasing laser power, enabling depths imaging of up to 100 µm (**Supplementary Fig. 8**).

We next assessed the suitability of small biopsies in the 3D-NaissI method for immunostaining. Due to inherent random sectioning and the large size of CMs, the fresh biopsies comprise both intact and sectioned CMs. Thus, we hypothesized that, without permeabilization, a selective antibody targeting an intracellular CM protein could label sectioned CMs while leaving intact CMs unlabeled (**Fig. 4a**). In agreement, fresh biopsies incubated with an anti-α-actinin antibody for 48 hours followed by a 48 hour-incubation with a fluorescent complementary secondary antibody, showed specific α-actinin staining in sectioned CMs, while intact CMs remained unlabeled (**Fig. 4b**). In contrast, PFA-fixed biopsies exhibited positive α-actinin staining in all CMs (**Fig. 4c**), agreeing with PFA-induced permeabilization. Saponin permeabilization allowed antibody penetration into intact cells without significant tissue disruption (**Supplementary Fig. 9a**), as demonstrated by similar measurements of the CM inter-lateral space (**Supplementary Fig. 9b**). Immunofluorescence imaging depth also linearly increased with extended incubation time, but optimal staining depth did not exceed 20 µm compared to small molecule staining (**Supplementary Fig. 10a**), likely influenced by antibody size. It is worth noting that the duration for antibody-based fluorescent staining can be streamlined to just 48 hours prior to imaging (**Supplementary Fig. 10b**). These depth variations were primarily influenced by fluorescent molecule size rather than laser penetration properties, as they persisted under similar laser parameters (**Supplementary Fig. 11**). Notably, immunofluorescent labeling on fresh tissue demonstrated remarkable selectivity for most tested antibodies, regardless of the manufacturer, thus supporting the hypothesis of frequent epitope masking in tissues when using fixatives.

**Figure 4.**
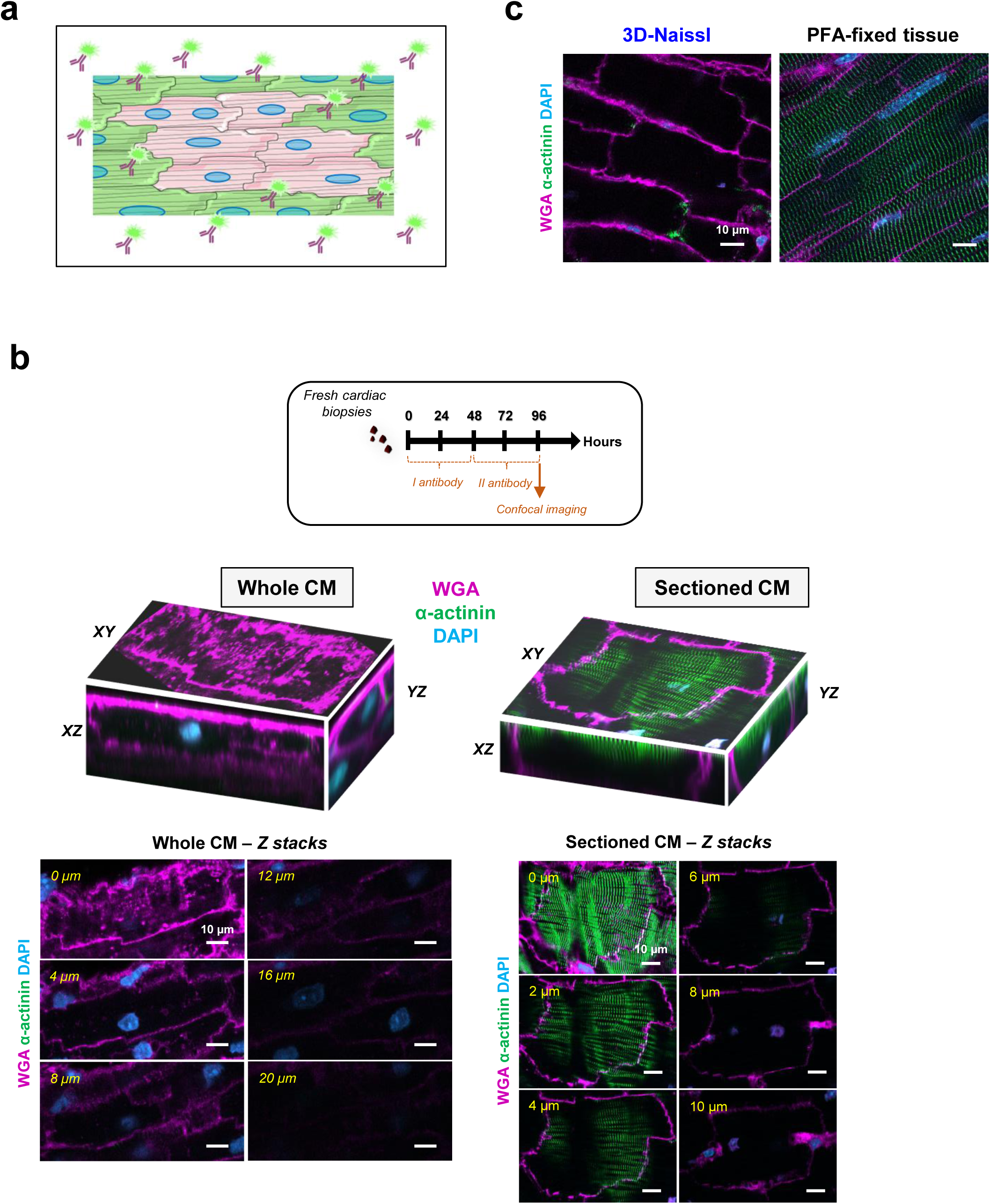
Immunofluorescent staining of fresh cardiac biopsies using 3D-NaissI method. **a** Schematic illustrating the hypothetical antibody penetrance in fresh cardiac biopsies, highlighting staining differences between whole and transected cardiomyocytes. **b.** Representative images (3 independent experiments) display XY, XZ, YZ projections and associated Z-stacks, illustrating fluorescent staining of a whole cardiomyocyte (left panel) or a sectioned cardiomyocyte (right panel) for wheat germ agglutinin (WGA), α-actinin, DAPI in fresh cardiac biopsies (3D-NaissI, 2-month old male mice). Both cases exhibit WGA-rod shape surface staining, with a notable absence of α-actinin staining is observed exclusively in the whole cardiomyocyte. **c** Representative images (3 independent experiments) of fluorescent staining for WGA, α-actinin, DAPI in the cardiac tissue from 2-month-old male mice, highlighting the contrast in antibody penetrance between fresh biopsies (3D-NAissI) with whole cardiomyocytes and PFA-fixed-tissue with permeabilized cardiomyocytes.

### Native expression of cardiac proteins using 3D-NaissI method: advancing scientific comprehension and interpretations

To assess the efficacy of the 3D-NaissI method, we explored the native expression patterns of both well-known and less-known cardiac proteins in fresh biopsies (myocardium) from left ventricles of mice.

We first examined connexin-43 (Cx43), a pivotal transmembrane protein in CM gap junctions essential for heart conduction, comparing conventional imaging of FFPE-cardiac tissues and the 3D-NaissI method. The imaging of FFPE-cardiac tissue confirmed the expected Cx43 localization at intercalated discs in control mice (**Fig. 5a**). Intriguingly, the 3D-NaissI method unveiled an additional presence of Cx43 at the CM lateral membrane (**Fig. 5b**), resembling the documented "Cx43 lateralization" observed in injured hearts[17–19]. To comprehensively understand these findings, we extended our analysis to cardiac tissue from X-linked muscular dystrophy (MDX) mouse model, a model of Duchenne muscular dystrophy showcasing "Cx43 lateralization" [20, 21]. As anticipated, Cx43 lateralization was evident in both FFPE (**Fig. 5a**, **MDX**) and fresh cardiac tissue (**Fig. 5b**, **MDX**) from adult MDX mice. However, in fresh biopsies, MDX mice exhibited distinct Cx43 expression patterns, characterized by a diffuse and punctuated distribution around the CM surface, contrasting with large clusters in control mice (**Fig. 5b**). While these results confirm issues with Cx43 expression/localization in the MDX mouse model, they demonstrate that that Cx43 lateralization is not pathology-specific but a hallmark of normal cardiac tissue in adult mice.

**Figure 5.**
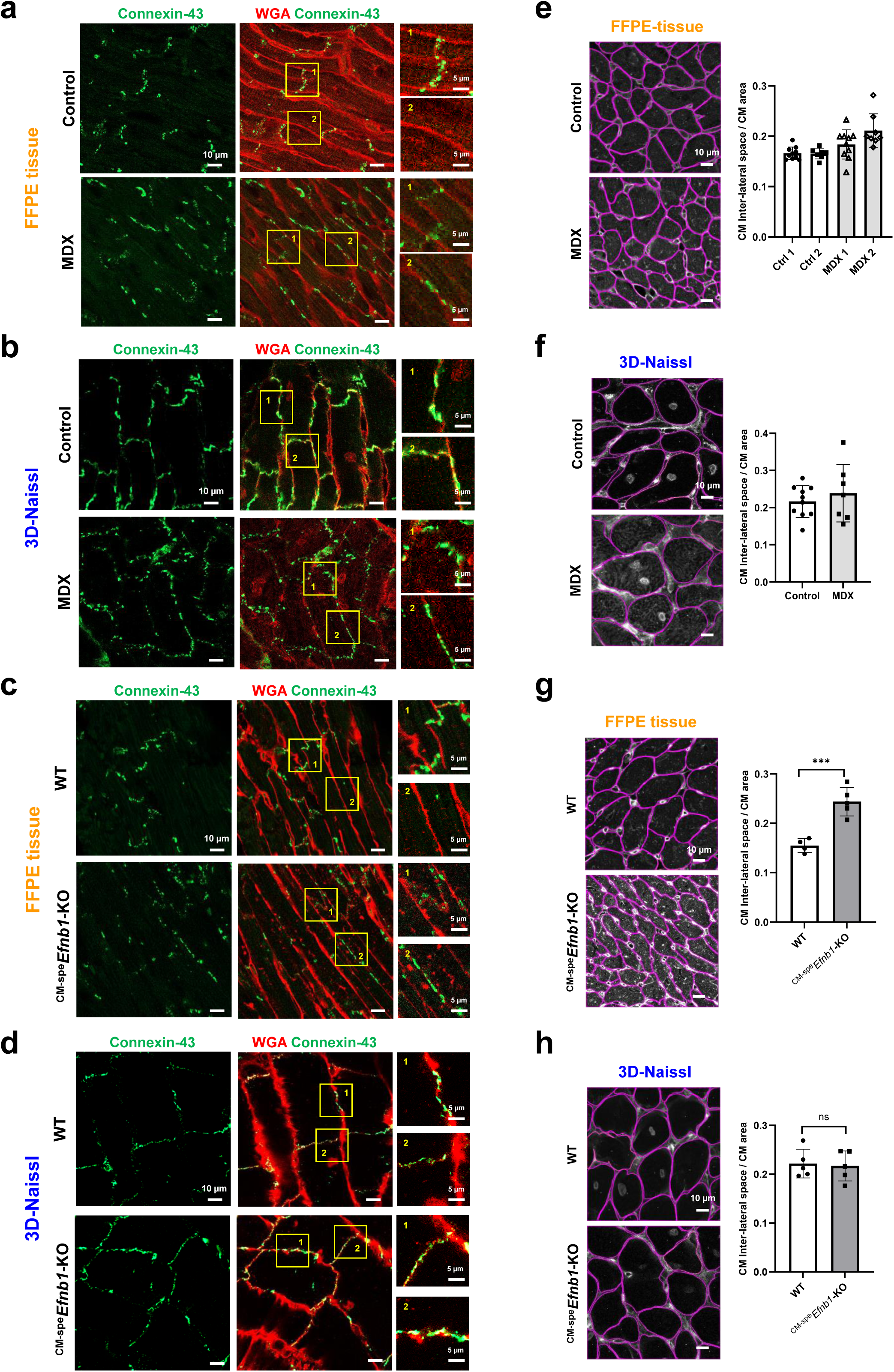
Artifacts of “connexin 43 lateralization” in formalin-fixed paraffin-embedded (FFPE)-cardiac tissues revealed through 3D-NaissI. **a-d** Representative images of connexin-43 immunostaining in the presence of fluorescent wheat germ agglutinin (WGA) in fresh-(3D-NaissI) or FFPE-cardiac tissue from 2-month old male X-linked muscular dystrophy (MDX) mice and their respective controls (C57BL6J OlaHsd) or WT and cardiomyocyte-specific *Efnb1* KO mice (CM-spe*Efnb1*-KO). Zoomed-in images (right panels) illustrate the localization of connexin-43 at the lateral membrane or the intercalated disc of the cardiomyocyte. **e-h** Corresponding quantification of the inter-cardiomyocyte (CM) space in fluorescent-WGA-stained biopsies from a-d d. Data are mean ± s.d. (**e**) *n*=2 mice /group; (**f**) *n*=1 mice/group; (**g, h**) *n*=4-5 mice/group (5-10 biopsies/mouse, 1 image /biopsy), unpaired Student’s *t*-test.

The absence of Cx43 immunostaining at the CM lateral membrane in FFPE-control mice may result from epitope masking, as it commonly occurs with this technical procedure[22], or restricted antibody access. This limitation may result from CM shrinkage and reduced lateral space between CMs induced by formalin fixation, as previously demonstrated (**Fig. 3b**). Accordingly, in MDX mice, the trend towards increased intercellular space in FFPE tissues (≥0.2; **Fig. 5e**) compared to control mice (˂ 0.2) supported improved antibody access, facilitating the detection of Cx43 lateral staining. Further supporting this hypothesis, the increased intercellular space in MDX-FFPE tissues closely approached that observed in the 3D-NaissI method (**Fig. 5f**). These findings were substantiated in CM-specific *Efnb1* KO mice, a model of tissue cohesion loss that we have previously described[23, 24]. While the 3D-NaissI method demonstrated similar CM inter-lateral space in both wild-type (WT) and KO mice (**Fig. 5h**), a significant increase was measured in FFPE-treated tissues from KO mice (**Fig. 5g**). Similar disparities between FFPE tissue and fresh tissue were noted when quantifying CM area (**Supplementary Fig. 12**), further substantiating the phenomenon of CM shrinking under FFPE conditions. Consistent with the expanded inter-cellular space observed in FFPE tissues from KO mice, we observed Cx43-specific staining at both the intercalated disk and the lateral membrane in the KO mice, while a unique staining pattern in the intercalated disk was depicted in WT mice (**Fig. 5c**). In contrast, the 3D-NaissI method unveiled Cx43 staining at both the CM lateral membrane (large patches) and the intercalated disc in control WT mice (**Fig. 5d**). Notably, in KO mice, a more continuous labeling was observed all around the CM surface (**Fig. 5d**), contrasting with our initial description in cryo-fixed tissue[23].

In summary, the 3D-NaissI method distinctly revealed native Cx43 expression in both the lateral membrane and the intercalated disc of healthy adult CMs *in situ*, challenging the conventional characterization of "Cx43 lateralization" as pathology-specific. The “Cx43 lateralization”, specifically described in conventional FFPE-pathological cardiac tissues and not in normal tissues, more likely reflects a technical artifacts leading to tissue shrinking with a decreased CM area and increased CM inter-lateral space. Now, whether the pathological phenotype of the CM interlateral space or the CM area observed in FFPE-treated tissue is a technical idiosyncrasy or an exacerbation of the pathological phenotype remains an open question.

Beyond the challenges encountered in visualizing proteins like Cx43 at the lateral side of CMs using conventional FFPE-processed heart sections, numerous transmembrane proteins remain difficult to image at the CM lateral membrane within the tissue context. Claudin-5, a tight junctional protein atypically localized at the CM lateral membrane[23, 25], exemplifies this challenge and has proven elusive in FFPE-cardiac tissue despite various protocols and antibodies, as illustrated in **Fig. 6a, b**. In contrast, the 3D-NaissI procedure applied to fresh cardiac biopsies allowed clear visualization of specific Claudin-5 staining on the inner face of the CM surface using an antibody targeting an intracellular epitope (**Fig. 6a**), while specific staining of the outer CM surface was achieved using an antibody directed against an extracellular epitope (**Fig. 6b**). Again, an additional advantage observed when utilizing fresh cardiac biopsies for fluorescent immunostaining with the 3D-NaissI method was the absence of nonspecific fluorescence. The effective visualization of transmembrane proteins, such as Claudin-5, at the CM lateral membrane within the tissue context demonstrates the superiority of the 3D-NaissI approach over conventional FFPE processing.

**Figure 6.**
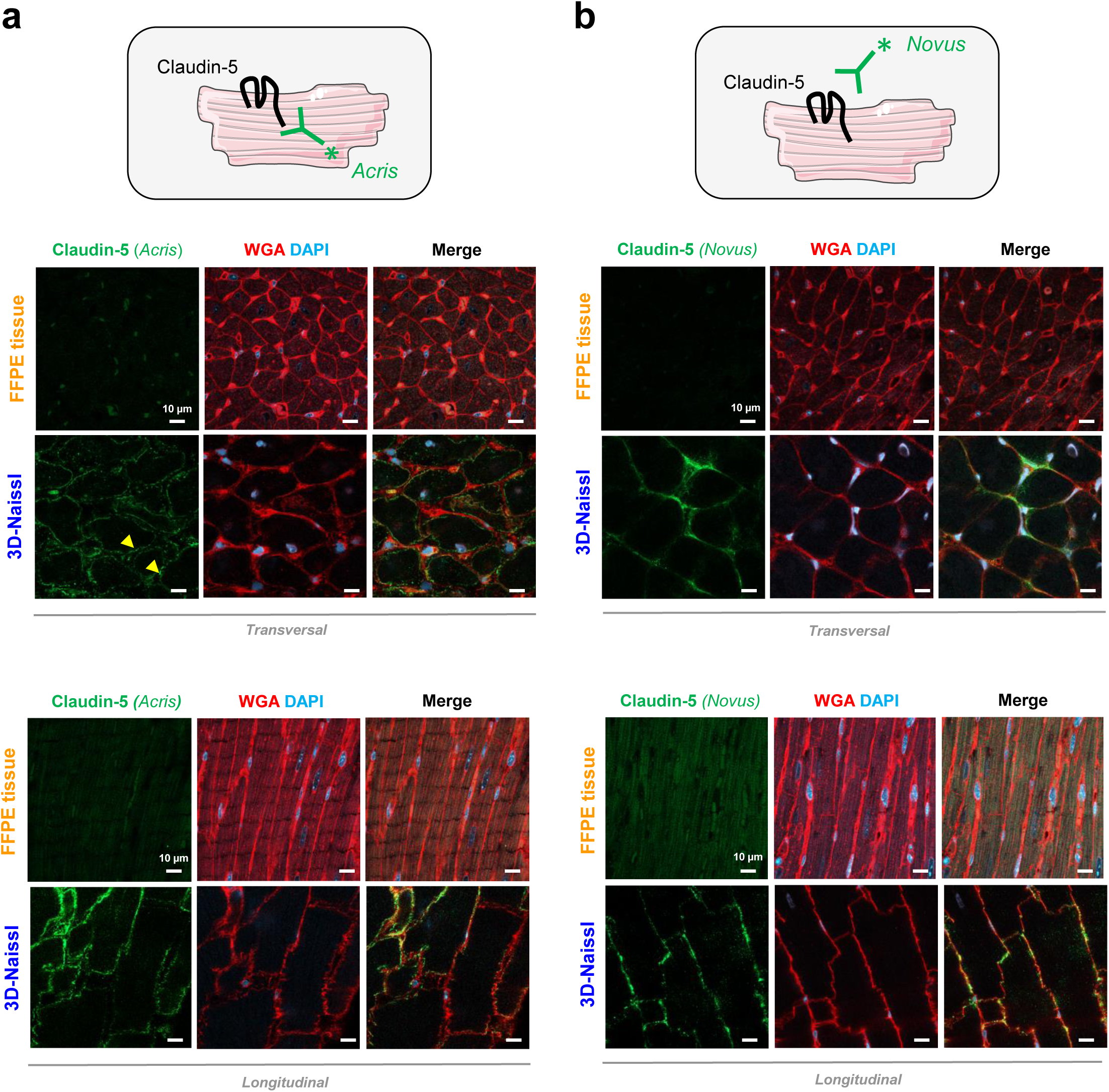
Visualization of the native expression pattern of Claudin-5 using the 3D-NaissI method. Representative confocal images (3 independent experiments) of Claudin-5 immunostaining in both fresh (3D-NaissI) or formalin-fixed paraffin-embedded (FFPE) cardiac tissue from 2-month-old male mice, in the presence of wheat germ agglutinin (WGA) and DAPI (*upper panels*: transversal view; *lower panels*: longitudinal view). Antibodies targeting either an intracellular epitope (**a**, Acris antibody) or an extracellular epitope (**b**, Novus antibody) of Claudin-5 were used.

The need for detailed mapping of protein expression within human cardiac tissues becomes evident in understanding specific cardiac diseases, particularly in the context of the recent COVID-19 pandemic wherein patients may experience severe and long-term cardiac complications that remain not fully understood[26]. In this critical context, the mapping of angiotensin-converting enzyme 2 (ACE2) expression, the receptor for SARS-CoV-2, was crucial. While transcriptomic profiling of ACE2 in the human heart has been reported[27, 28], protein expression and spatial distribution within the tissue remain underexplored. Immunohistochemistry, used in relatively few studies, has yielded highly variable results and poor resolution quality[29–32]. Here, we harnessed the 3D-NaissI method to map ACE2 spatial expression in fresh myocardial tissues from adult mice, rats, and humans, using a knockout-validated anti-ACE2 antibody. Similar ACE2 expression patterns were observed in all examined cardiac tissue species (**Fig. 7a**), including specific expression in vascular cells confirmed by co-localization with Iso-B4 (**Fig. 7b**), as classically reported. Notably, ACE2 was detected at the plasma membrane of CMs exhibiting co-localization with caveolin-3 on the surface and within intracellular T-tubule invaginations (**Fig.7c**). Significant ACE2 expression was also noted in the CM nuclei (**Supplementary Fig. 13a**). The specificity of ACE2 expression in CMs was further confirmed by western-blot analysis using lysates from adult primary CMs purified from mouse hearts (**Fig.7d**). In contrast, specific ACE2 expression in CMs was elusive using FFPE-tissues following supplier-recommended protocols, being detected only in vessels (IsoB4 co-localization; **Supplementary Fig. 13b**). The distinctive patterns of ACE2 expression in CMs, highlighted through the 3D-NAissI method, reveal potential mechanisms underlying SARS-CoV-2 infection and its impact on cardiac pathophysiology, particularly in COVID-19 cardiomyopathies. Additionally, the identification of ACE2 in the nuclei of CMs unveils a novel potential target for the SARS-CoV-2 spike protein, suggesting its involvement in dysregulation of nuclear signaling and contributing to the pathogenicity of SARS-CoV-2 in the heart. Accordingly, a recent study has revealed the presence of a nuclear localization signal sequence within the spike protein, indicating its putative potential for nucleus translocation[33].

**Figure 7.**
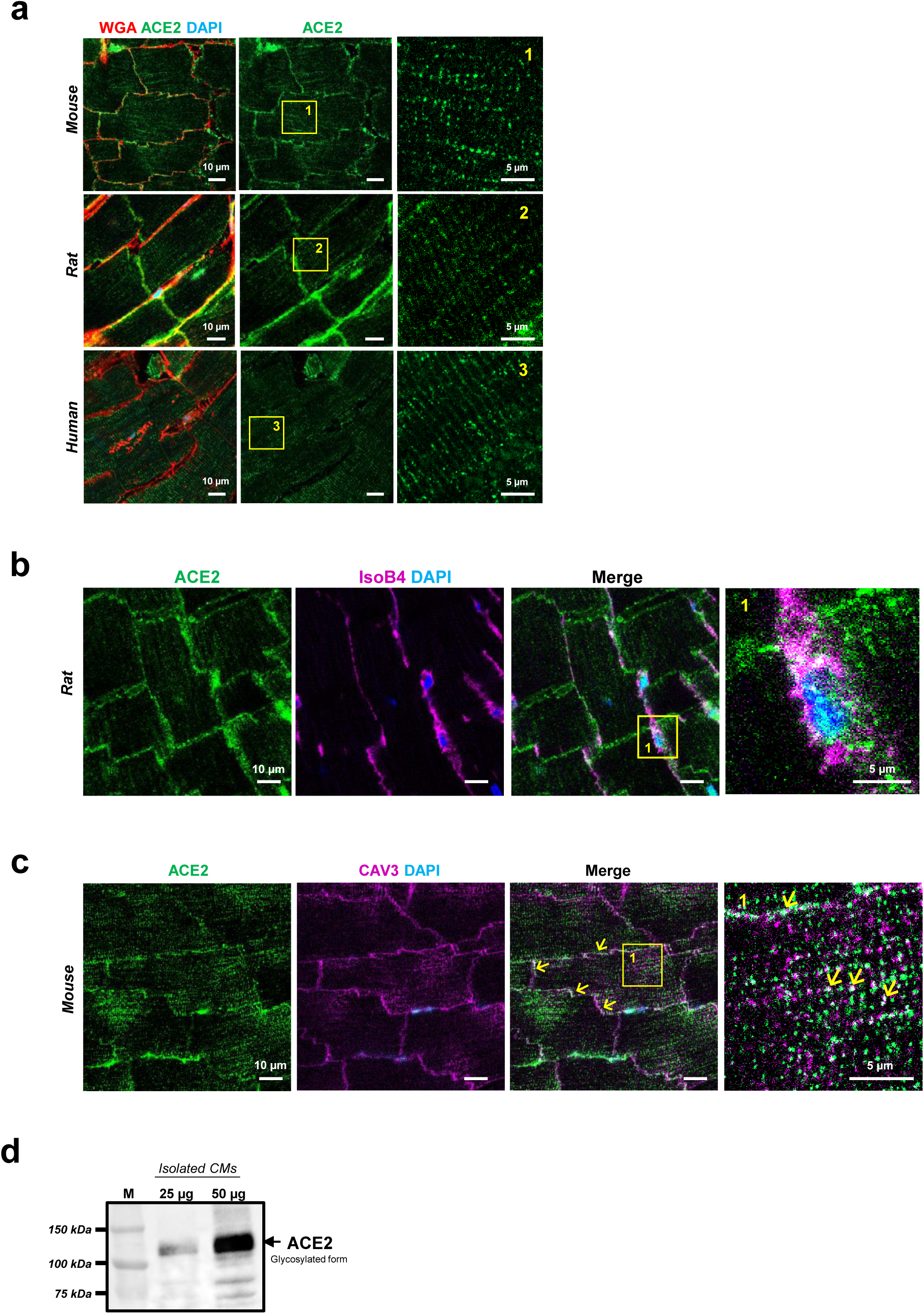
Visualization of native ACE2 expression in cardiac tissue though the advanced 3D-NaissI Method. Representative images (3-5 independent experiments) of: **a** ACE2 immunostaining with fluorescent wheat germ agglutinin (WGA) and DAPI in fresh cardiac tissue (3D-NaissI) from 2-month-old male mice, rats or 19-year-old-man. Zoomed-in images (right panels) highlight ACE2 localization within cardiomyocytes. **b** ACE2-Isolectin B4 (IsoB4)-DAPI co-staining in fresh cardiac tissue from 2-month old male rats. Zoomed-in image (right panels) highlights ACE2 localization within vascular cells (Iso-B4 positive cells). **c** ACE2-Caveolin-3 (CAV3)-DAPI co-staining in fresh cardiac tissue from 2-month old male mice. Zoomed-in image (right panels) emphasize ACE2 positive co-localization with Caveolin-3 within cardiomyocytes. **d** Representative ACE2 expression quantified by western blot in 25 and 50 µg lysates from purified adult cardiomyocytes from 2-month-old male mouse.

A contemporary challenge in cardiology revolves around comprehending the cardiac benefits associated with Sodium-glucose cotransporter 2 inhibitors (SGLT2i) therapeutic interventions through the examination of SGLT2 protein expression patterns in the heart. Recently, iSGLT2 have shown promise in reducing cardiovascular death and hospitalization in patients with Heart Failure with Preserved Ejection Fraction (HFpEF)[34], which accounts for over 50% of heart failure (HF) cases. Despite *in vitro* studies demonstrating direct cardiobenefits of SGLT2 inhibitors[35, 36], this finding contradicts the prevailing belief that SGLT2 expression is absent in both healthy and failing myocardium[37], despite conflicting results in the literature[38]. To address this discrepancy, our investigation focused on discerning the expression profile of SGTL2 in fresh myocardial biopsies obtained from adult mice, using the 3D-NaissI method. Our results unequivocally revealed specific punctate and lamellar labeling of SGLT2 within CMs and below the CM plasma membrane (**Fig. 8a**). This staining did not co-localize with caveolin-3 (**Fig. 8b and Supplementary Fig. 14a**), excluding the expression of SGTL2 on the plasma membrane side of the CM. However, co-localization was observed with sarcoplasmic protein Calsequestrin (CASQ, **Fig. 8c and Supplementary Fig. 14b**), providing substantial evidence for specific SGTL2 expression within the sarcoplasmic reticulum of the CM. Supporting SGTL2 expression in the sarcoplasmic reticulum, empagliflozin was shown to enhance contractility and Ca^2+^ transients in isolated CMs[36]. This specific expression of SGTL2 was further confirmed through western-blot analysis using isolated adult CMs (**Fig. 8d**). In contrast, we failed to detect SGTL2 in FFPE-tissue following supplier-recommended protocol (**Supplementary Fig. 14c)**. The identified specific expression of SGLT2 in the sarcoplasmic reticulum of CMs may have implications for the regulation of Ca^2+^ uptake to this compartment during diastole. This novel finding offers a compelling and alternative explanation for the observed therapeutic benefits of iSGLT2 in HFpEF patients.

**Figure 8.**
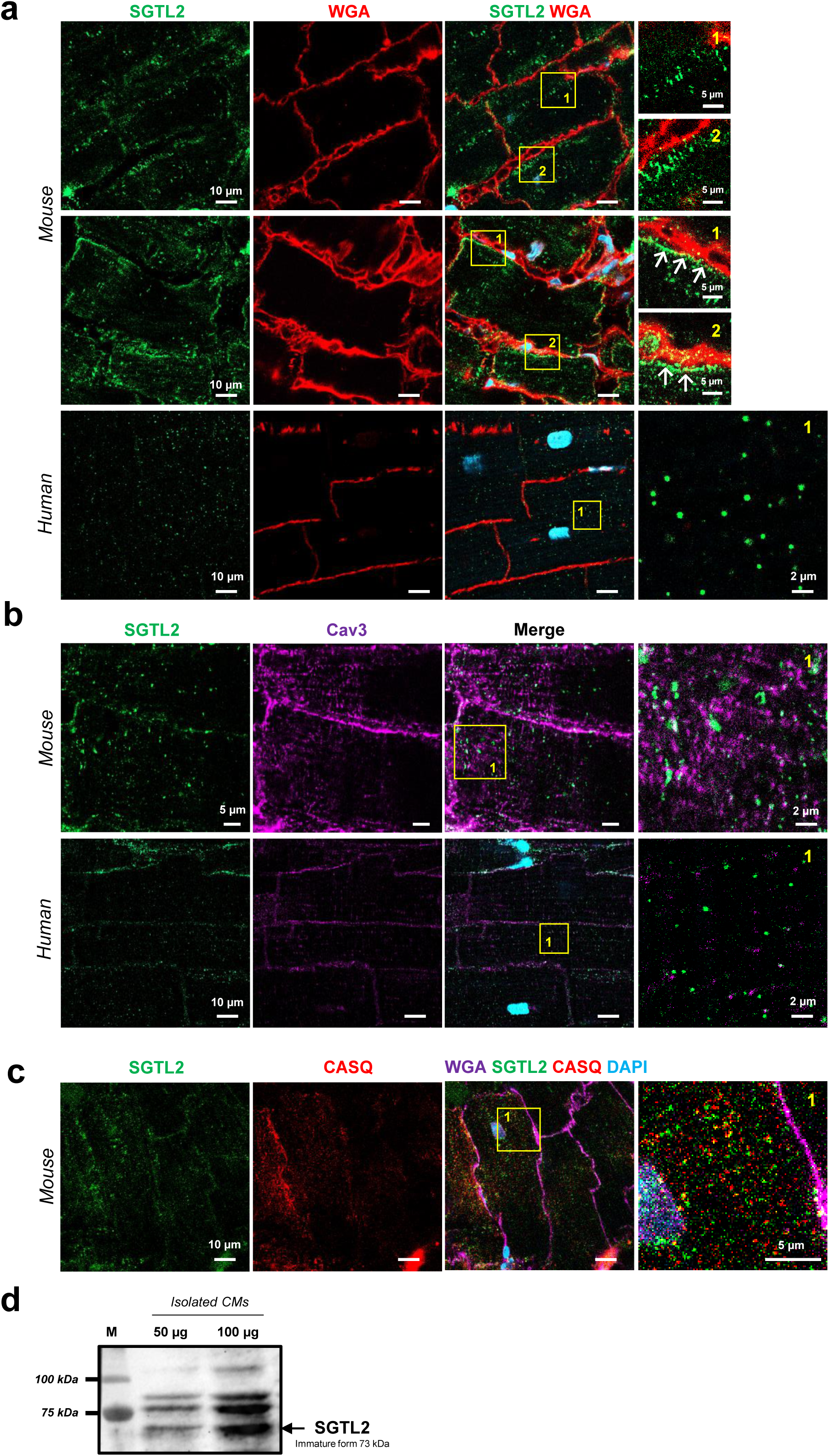
Visualization of native SGTL2 expression in cardiac tissue though the advanced 3D-NaissI Method. Representative images (3-4 independent experiments) of: **a** SGTL2 immunostaining with fluorescent wheat germ agglutinin (WGA) and DAPI in fresh cardiac tissue (3D-NaissI) from 2-month-old male mice (3 independent experiments) or a 20-year-old woman. Zoomed-in images (right panels) highlight the punctuated and laminar SGTL2 staining within cardiomyocytes and beneath the plasma membrane (WGA-positive). **b** SGTL2-Caveolin-3 (CAV3) co-staining in fresh cardiac tissue from 2-month-old male mouse or a 20-year-old woman. Zoomed-in image (right panels) emphasizes the absence of SGTL2 co-staining with Caveolin-3. **c** SGTL2-Calsequestrin (CASQ)-WGA-DAPI co-staining in fresh cardiac tissue from 2-month-old male mice. Zoomed-in image (right panels) emphasizes SGTL2 positive co-localization with Calsequestrin (yellow dots) within cardiomyocytes. **d** Representative SGTL2 expression quantified by western blot in 50 and 100 µg lysates from purified adult cardiomyocytes from 2-month-old male mouse.

In addition to unraveling the expression of elusive cardiac proteins, the 3D-NaissI method facilitates the assessment of native expression / organization of known proteins within the natural context of cardiac tissue. It proves instrumental in investigating the native organization of components integral to excitation-contraction coupling (E-C coupling) within cardiac tissues, a domain predominantly explored in isolated CMs[39]. Unlike isolated CMs in culture, CMs within the tissue experience longitudinal mechanical constraints at the intercalated disc junctions and lateral constraints through interactions with the extracellular matrix and surface crests neighbored CMs. Both of this factors may influence the expression and spatial organization of proteins involved in the regulation of the E-C coupling. We applied the 3D-NaissI method to visualize Junctophilin 2 (JPH2) in fresh cardiac biopsies, a crucial structural protein forming ’dyads’ that link the sarcoplasmic reticulum to the T-tubule membrane, in conjunction with the Ryanodine receptor (Ryr) located on the sarcoplasmic reticulum. These proteins play pivotal roles in E-C coupling and are prone to spatial disorganization in HF[40]. As depicted in **Supplementary Fig. 15a**, the expression of JHP2 in fresh myocardial samples from mice and humans aligns with the expected species differences in T-tubule organization, with smaller species exhibiting more intricate arrangements compared to larger mammals[40]. Consistent with the reduction of T-tubules observed during aging[41], JHP2 exhibited reduced expression and spatial organization in older patients compared to younger individuals (**Supplementary Fig. 15b**). The application of Airyscan technology allowed for the acquisition of JHP2 and Ryr expression patterns with exceptional resolution (**Supplementary Fig. 15c and Supplementary Movie 1**). In the future, the 3D-NaissI method holds promise for enhancing the analysis of E-C coupling component disorganization within a tissue context across diverse cardiac pathologies.

The 3D-NaissI method provides only a small window of tissue visibility compared to the observation of the entire heart, yet excels in revealing intricate 3D architecture within specific cardiac compartments. The 3D reconstruction of the left ventricle myocardium stained with fluorescent-WGA unveiled previously undocumented ring-shaped structures surrounding α-actinin positive-CMs along their longitudinal axis in the adult mouse heart, a pattern conserved in rats and humans (**Fig. 9a, b and Supplementary Fig.16a**). These structures, found throughout the myocardium, endocardium, and epicardium (**Supplementary Fig. 17**), were compromised following PFA-biopsy fixation (**Supplementary Fig.18a**). Close examination of WGA-staining revealed that the ring structures emanate from the cytoplasmic extensions of specific cells lying on the endothelial cell prolongations at the lateral surface of CMs, as indicated by nuclear staining in this location. These structures exhibited 100 % IsoB4 immunoreactivity (**Fig. 9c and Supplementary Fig.16b and 18b**), suggesting a vascular origin. Drawing parallels with ring-like structures formed by vascular smooth muscle cells (VSMCs) in the macrovasculature[42], we hypothesized that pericytes, with their remarkable extension and connectivity potential (https://doi.org/10.1007/s43152-021-00029-w), might mimic, on the microcirculation side and around the CM myofiber, the spatial organization of VSMCs. Supporting this hypothesis, the ring structures tested positive for the NG2 pericyte marker (**Fig. 9d and Supplementary Fig.16c**), and exhibited 100% positivity for laminin 2 (**Fig. 9e**), suggesting that cardiac pericytes play a crucial role in the production of the CM basement membrane, a function recently demonstrated in other tissues[43, 44]. Accordingly, a distinct spatial arrangement of laminin-2 receptors, α-dystroglycan (**Fig. 9f**) and β1 integrin (**Supplementary Fig. 18c**), was observed along the laminin 2 positive ring structures on the CM surface. This reveals a novel spatial organization of pericytes in cardiac tissue[45], forming 3D-ring structures encompassing the rod-shaped CMs. This unique organization challenges the conventional understanding of ring structures formed by VSMCs around macrovasculature or pericytes around microvasculature[42]. The substantial coverage of CM lateral surfaces by pericytes, as revealed in this study, underscores a new pivotal role of pericytes in CM function.

**Figure 9.**
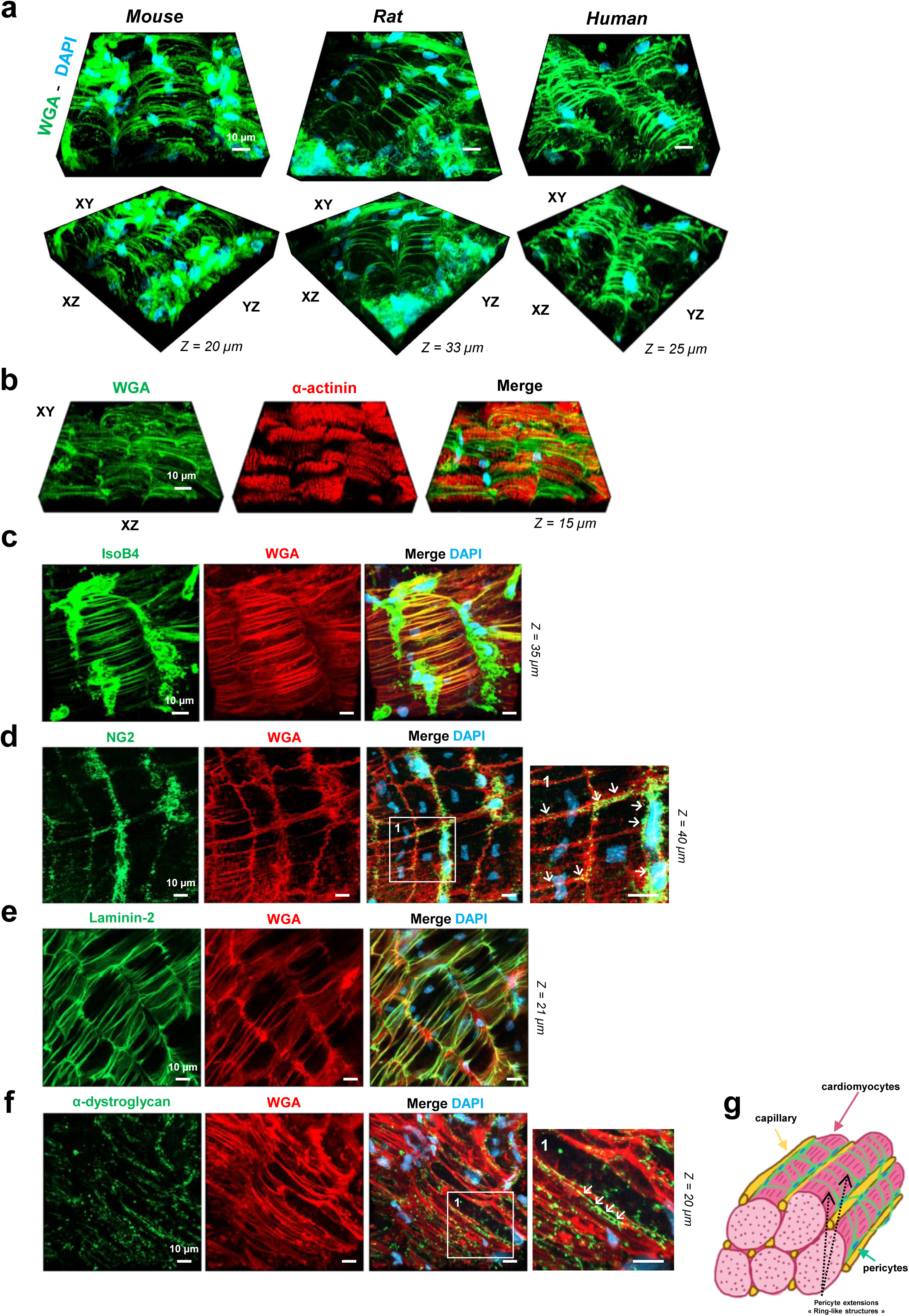
Unveiling a novel 3D architectural paradigm in fresh cardiac tissue through advanced 3D-NaissI imaging. Representative images three-dimensional confocal imaging (3-5 independent experiments) of: **a** Wheat germ agglutinin (WGA)-DAPI-stained fresh cardiac biopsies from 2-month-old male mice, rat or a 26-year-old-man, revealing unidentified WGA-positive tube-like structures. These structures intricately surround cardiomyocytes along their longitudinal axis, as depicted by α-actinin positive staining (**b**). These novel 3D-structures are: (**c**) Isolectin B4 (IsoB4)-positive, (**d**) Neural/glial antigen 2 (NG2)-positive, (**e**) Laminin-2-positive and (**f**) meticulously organized along the α-dystroglycan positive-staining at the surface of cardiomyocytes. **g** Schematic illustrating a novel model of pericyte architecture in the cardiac tissue.

While optimized for imaging native cardiac tissue, the 3D-NaissI method’s applicability extends to challenging tissues such as the lipid-enriched brain, solid-cohesive skin, spongy air-filled lung tissue, and the gut. The method was applied directly to 1 mm^3^ fresh biopsies extracted from the mouse cortex, human skin, mouse lung, or mouse gut, successfully enabling imaging of all these tissues (**Supplementary Fig. 19**).

## Discussion

This study introduces the 3D-NaissI method as an innovative and robust approach for imaging fresh native cardiac tissue biopsies, offering distinct advantages over traditional histological techniques. Our findings underscore the method’s exceptional ability to preserve tissue integrity in hypothermic conditions, thus establishing a reliable platform for short-term imaging studies. By preserving tissue integrity, the 3D-NaissI method enables detailed imaging of cellular and molecular architecture, facilitating a comprehensive examination of native cardiac protein expression. Several pivotal insights have emerged from our results, underscoring the far-reaching implications of the 3D-NaissI method for advancing cardiac research.

In contrast to conventional tissue techniques, such as light and electron microscopy, which typically involve the use of chemical cross-linking agents to preserve tissue integrity, the 3D-NaissI method is entirely fixative-free. This approach relies on the use of fresh tissue biopsies throughout the entire process of fluorescent immunostaining and confocal imaging, successfully circumventing major artifacts associated with chemical fixation:

i/ The 3D-NaissI method facilitates the preservation of antibody epitopes, mitigating issues like epitope masking, conformational changes in proteins, disruption of protein-protein interactions, even protein aggregation artifacts. As a result, the 3D-NaissI method enables the successful use of a wide range of commercially available antibodies described for immunofluorescence, which stands in contrast to classical immunohistological techniques that often require unmasking strategies or prove ineffective with many antibodies.
ii/ The 3D-NaissI method significantly reduces autofluorescent background signals and nonspecific antibody binding. This reduction contributes to a high signal-to-noise ratio, enabling the detection of low levels of protein expression without the need for additional blocking steps to minimize nonspecific signals.
iii/ The 3D-NaissI method excels in preserving the native cellular and molecular spatial organization. Hence, our study delves into the intricate architecture of cardiac tissue, uncovering previously undocumented ring-like structures formed by pericytes around CMs. This novel 3D organization emphasizes the significance of pericytes in cardiac tissue and challenges conventional notions of their spatial arrangement, which has been exclusively described relative to the vascular cells until now This intricate structure, lost after tissue fixation, underscores the method’s superiority. Unlike fixed biopsies, fresh samples retain the inter-lateral space between CMs, reflecting natural tissue cohesion. Fixation-induced shrinkage alters intercellular space, potentially masking epitopes. Notably, 3D-NaissI minimizes cell size changes seen in fixed conditions, preventing significant modifications to biomolecule organization.

Challenging prior assumptions, these findings underscore the importance of studying protein expression in the native tissue context. Applying 3D-NaissI to analyze cardiac proteins like Cx43, ACE2, and SGLT2 reveals novel insights into their distribution. Particularly relevant during the COVID-19 pandemic, mapping ACE2 expression sheds light on cardiac complications related to infectious diseases. The unexpected lateralization of Cx43, as revealed in native healthy cardiac tissues, has challenged our understanding of cardiac pathophysiology. Contrary to previous assumptions, this discovery points more towards concerns about tissue decompaction (increased inter-lateral space between CMs) than a pathological localization of Cx43 on the lateral face of the CM. The presence of Cx43 at the CM lateral membrane prompts questions about its specific role in this location, challenging conventional wisdom regarding its typical presence at the intercalated disc and its established role in rhythmology and arrhythmogenesis[18].

In summary, the fixative-free nature of the 3D-NaissI method offers significant advantages over traditional techniques. It not only effectively preserves tissue integrity and spatial organization but also mitigates artifacts, thus enhancing the overall quality of imaging and analysis for a more comprehensive understanding of native tissue characteristics.

Beyond fixative omission, the 3D-NaissI method offers a significant advantage by obviating delipidation-based permeabilization, a conventional hurdle in intracellular antibody penetration. Preserving cell membrane integrity unlocks the visualization of native membrane proteins, a feat conventionally more challenging. Moreover, this technique is characterized by its simplicity, swift application, compatibility with standard confocal microscopes, and cost-effectiveness. However, limitations include reliance on small tissue samples and restricted fields of view, positioning the 3D-NaissI method as a complementary tool that synergizes with other classical tissue imaging techniques for affirming native protein expression in specific organ sub-compartments.

The use of fresh cardiac tissue in cardiology is not new, with the emergence of living myocardial slices (LMS) in recent years, albeit presenting technical challenges[46]. Unlike isolated cell preparations, LMS serves as a physiologically relevant *ex vivo* organotypic model, preserving the native intercellular network. However, current developments have predominantly focused on functional studies, concentrating on mechanical and electrical investigations[47]. Although maintaining LMS functionality for extended periods involves specific *ex vivo* culture procedures[13, 46, 48], such as the use of thick cardiac slices (∼300 µm) and environmental factors like oxygenation, ionic culture medium, and electromechanical stimulation, its applicability for microscopic imaging is constrained. Typically, fixatives are necessary, compromising the tissue’s native properties. In this context, the 3D-NaissI method emerges as a complementary approach to LMS, facilitating the short-term utilization of fresh cardiac biopsies specifically for imaging purposes.

In conclusion, the 3D-NaissI method stands out as a powerful tool for short-term imaging of fresh native cardiac tissue biopsies, offering relative high preservation of tissue integrity, viability, and native protein expression. Its versatility is evident not only in cardiac research but also in imaging various tissues, showcasing its potential applications. The method’s ability to provide detailed insights into the 3D organization of cellular components and protein expression patterns makes it a valuable addition to the toolbox of researchers investigating biological systems in their native state. Hence, the 3D-NaissI method holds great promise for both research and clinical applications. Beyond its potential to complement existing databases like the Human Protein Atlas (www.proteinatlas.org), the method distinguishes itself with clinical diagnostic utility. For instance, the use of fresh cardiac biopsies obtained during surgeries could yield valuable insights into cellular and molecular aspects of cardiac diseases, extending its applicability to other medical fields such as oncology. Integrating this methodology into surgical procedures might not only enhance our understanding of diseases but also facilitate translational research, bridging the gap between basic research and clinical practice. In line with the clinical translational aspect, the ability to image fresh tissue biopsies derived from human autopsy samples adds a significant dimension to the 3D-NaissI method. This provides access to a larger tissue collection, including control tissues that are often scarce in surgical contexts. However, it is essential to note that the imaging results of autopsy biopsies are contingent upon the clinical context of the patient. Overall, the 3D-NaissI method emerges as a versatile tool with profound implications for advancing biomedical knowledge and improving patient care.

## Methods

Further material and methods details used in his manuscript are available on the Supplementary material online.

### Ethical statement

Procedures involving human cardiac samples were conducted at the Department of Forensic Medicine, Centre Hospitalier Universitaire de Toulouse (University of Toulouse, France), adhering strictly to the principles outlined in the *Declaration of Helsinki*. Furthermore, these procedures received approval in accordance with French legislation from the Agence de Biomédecine under registration number PFS21-015, with explicit consent obtained from the relatives, signifying their non-objection to the sampling process.

All animal experiments were performed in accordance with the European directive for the protection of animals usedfor scientific purpose and were approved of the French CEEA-122 ethical committee (CEEA 122 2015-28).

### Animal models and euthanasia

All studies were performed on 2-month old male C57BL/6J OlaHsd mice, Spague Dawley rats (purchased from Envigo, Huntingdon, United-Kingdom). Hearts from male X-linked muscular dystrophy mice (C57BL/10ScSnDmdmdx/J) designed as MDX, were obtained from Dr O. Cazorla, (PhyMedExp, Université de Montpellier, France) and from a colony maintained in their local animal facility (Plateau Central d’Elevage et d’Archivage, Montpellier, France). Two-month old male C57BL/6J OlaHsd mice purchased from Envigo were designed as control for the MDX mice. Cardiomyocyte-specific *efnb1* knock-out mice (KO) and their WT counterparts have already been described[23], and all studies using these mice were performed on male and age-matched littermate mice.

All animals were maintained in the animal facility at the UMS06-Centre régional d’exploration fonctionnelle et de resources expérimentales-CREFRE (Toulouse, France) under specific-pathogen free (SPF) conditions and following institutional animal use and care guidelines. Animals were housed conventionally in controlled humidity and temperature, operating on12 h of light and dark cycle with food and water available ad libitum. Animals were anesthetized with an intraperitoneal (i.p.) injection of 50 mg/kg Dolethal (Vetoquinol, Lure, France) in the presence of 0.1 mg/kg buprenorphine and euthanasia was performed after chest opening by promptly excising the beating heart to prepare fresh cardiac biopsies.

### Preparation of fresh cardiac biopsies

For mice and rats, immediately after removal, the beating heart was transferred in cold PBS and gently pressed to squeeze blood out. Then, a cross-sectional heart slice (∼ 3 mm thick) was briefly washed in 3 successive cold PBS baths. The left ventricle was excised, the endocardium and epicardium were excised and the remaining myocardium was then cautiously sliced (∼1 mm^3^ biopsies) on a glass slide on ice using a very fine and sharp scalpel. Similar procedure was performed for the human cardiac biopsies, except that only a small portion of the left ventricular myocardium was harvested from autopsy hearts.

### Fluorescent staining of fresh cardiac biopsies

#### Tissue integrity study

Fresh biopsies were incubated in 500 µL PBS, HBSS or cardioplegic solution in 24-well plates at 4°C, 37°C or room temperature under constant rotational agitation for 2 days in the dark in the presence of Alexa Fluor-WGA, DAPI and Alexa Fluor-IsoB4. For the kinetics study, fresh biopsies were incubated in cold PBS for 0, 2, 4, 7 or 10 days and stained with Alexa Fluor-WGA and DAPI for 4 hours before confocal image acquisition.

#### Tissue viability study

Fresh biopsies were incubated in 500 µL PBS, HBSS or cardioplegic solution 24-well plates at 4°C, 37°C or room temperature under constant rotational agitation for 2 days in the dark and in the presence of Alexa Fluo-WGA, DAPI and MitoTracher®Red CMXRos. For the kinetics study, fresh biopsies were prepared on Day 0 and subjected to 2 days of staining in PBS at 4°C in the dark with MitoTracher®Red CMXRos, Alexa Fluor^TM^-WGA and DAPI before confocal imaging at day 2, 3, 4, 5 and 8 (**Supplementary Fig. 5a**). Alternatively, staining was performed in the presence of Di-8-ANEPPS and DAPI for 4 hours before confocal imaging at day 2, 4, 7 and 10 (**Supplementary Fig. 5b**).

#### Tissue immunostaining

Fresh biopsies were incubated in 500 µL PBS in 24-well plates at 4°C under constant rotational agitation in the dark with the primary antibody for 2 days, followed by a rapid washing step with cold PBS, incubation with the secondary antibody for 2 more days at 4°C under constant rotational agitation in the dark, and finally a prompt was with cold PBS before confocal imaging.

### 2D quantification of cardiomyocyte area and inter-lateral space

Quantification of cardiomyocyte (CM) area was performed on confocal images of paraffin-embedded heart sections (transverse sections) or fresh cardiac biopsies, both stained with Alexa Fluor^TM^488-WGA, enabling precise delineation of the CM surface. The CM areas were quantitatively assessed using Fiji (Image J, NIH) by manually tracing cellular contours from confocal microscope images (Zeiss LSM 900 confocal microscope, Carl Zeiss) captured at a magnification of 63X. Approximately 5-10 images per mouse were acquired to ensure appropriate statistical analysis. On the same 2D confocal images, inter-lateral space between CMs was quantified by subtracting the total CM area from the entire image area. To account for potential variations in CM area associated with cardiac phenotype, the inter-lateral space was finally quantified relative to the CM area and thus expressed as a ratio.

### Confocal microscopy and image acquisition/analysis

Fresh biopsies were mounted on concave microscope glass slides (Hecht Karl™, 42412010) filled with 80 µL cold PBS and covered with coverslips (Menzel-Gläser, MENZCB00190RA120) fixed with a silicon seal and immediately imaged.

Image acquisition was conducted on a ZEISS LSM 900 Axio Observer.Z1/7 with Airyscan 2 inverted confocal microscope using a X63 1.4 NA oil objective at a maximum resolution of 1024x1024 pixels. Three-dimensional stacks were acquired with a 1 µm z-spacing and x and y pixel size of 101.41 µm. The images presented in this paper represent the most representative selection from multiple acquisitions. Image visualization was performed using Zen Blue (Carl Zeiss, Germany) software. Three-D rendering images was carried out using Zen Blue and Imaris (Bitplane, Belfast, UK).

### Quantification of cAMP production in fresh cardiac biopsies

Quantification of intracellular cAMP was performed using the HTRF cAMP competitive immunoassay (*cAMP Gi kit, 62AM9PEB, Revvity*) according to the manufacturer’s instructions. Fresh mouse cardiac biopsies (1 biopsy ∼1-3 mg/well) were distributed in a 96-well white microplate (OptiPlate-96; PerkinElmer) and incubated or not (basal) with isoproterenol (10 µM) or carvedilol (10 µM) alone or in combination for 30 min at room temperature and in the presence of 0.5 mM IBMX to prevent phosphodiesterase-mediated cAMP degradation. After the addition of cryptate-labeled cAMP (donor) and anti-cAMP-d2 (acceptor) for 1 h, the specific FRET signals were calculated by the fluorescence ratio of the acceptor and donor emission signal (665/620 nm) collected using a modified Infinite F500 (Tecan Group Ltd). Conversion of the HTRF ratio of each sample into cAMP concentrations was performed on the basis of a standard curve to determine the linear dynamic range of the assay.

### Data Analysis-Statistics-Reproducibility

The n number for each experiment and analysis is stated in each figure legend. An unpaired Student t-test for parametric variables was used to compare two groups. One-way ANOVA with Tukey post-hoc test was used for multiple group comparisons (> 2 groups). The level of significance was assigned to statistics in accordance with their p values: p ≤ 0.05 *; p ≤ 0.01 **; p ≤ 0.001 ***; p ≤ 0.0001 ****). All graphs and statistics were generated using v9.2.0 (GraphPad Inc, San Diego, California).

## Declarations

## Acknowledgements

We thank support from Remy Flores-Flores at the PHI-histology imaging platform (I2MC-Toulouse, France) and the We-Met functional biochemistry platform from TRI Genotoul network facilities. We also thank Dr Olivier Cazorla from PhyMedExp (Université de Montpellier, France) for providing the mdx mice. Dr Corinne Lorenzo is acknowledged for helpful discussions on biopsy imaging, and we thank everyone from the ANR-18-CE06-0027-04 consortium for all their support in this project.

## Author contributions

C.G. conceived the idea and assisted N.P in realizing all the imaging experiments. N.P. performed all the small tissue biopsies required for the 3D-NaissI method, as well as the entire processing of the biopsies up to image acquisition. C.G.F. provided us with the human biopsies. C.G.F. and J.M.S. handled ethical authorization at the “Agence de Biomédecine” for the use of human biopsies obtained from autopsies for scientific purposes. V.P. conducted the functional studies (cAMP production) of the biopsies used in the NaissI method. A. W. conducted and quantified all Western blot experiments, as well as some staining experiments using the NaissI method. C.G wrote the paper with the help of C.K., J.M.S and CGF. All authors contributed to proofreading.

## Data Availability

The datasets generated during and/or analyzed during the current study are available from the corresponding author on reasonable request.

## Ethics approval

All animal experiments were performed in accordance with the European directive for the protection of animals used for scientific purpose and were approved of the French CEEA-122 ethical committee (CEEA 122 2015-28).

## Consent for publication

All the authors have approved and agreed to publish this manuscript.

## SUPPLEMENTARY METHODS

### Reagentsh

The following reagents were used in this study: Dulbecco’s Phosphate Buffered Saline (PBS) (Sigma-Aldrich, #D1408), Hank’s Balanced Salt Solution (HBSS) (Sigma-Aldrich, #H8264), Celsior® (Genzyme Corp., Boston, MA) cardioplegic solution for arrest and storage in heart transplantation, Paraformaldehyde aqueous solution (PFA) (Electron Microscopy Sciences, # 15714), Saponin (Sigma-Aldrich, #84510).

### Preparation of PFA fixed-cardiac biopsies

Fresh cardiac biopsies were immediately incubated in a 4 % Paraformaldehyde fixative solution for 1h at room temperature before fluorescent staining.

### Preparation of Formalin-Fixed Paraffin-Embedded (FFPE) heart sections

Hearts were excised, immediately fixed in 4% formol for 48h and finally embedded in paraffin and sectioned (transverse sections) at 5 μm intervals.

### Antibodies and probes for immunofluorescence studies

The unconjugated primary antibodies in this study were as follows: Rabbit monoclonal anti-ACE2 [EPR4435(2)] (Abcam, ab1082521, 1:50); Rabbit polyclonal anti-Junctophilin-2 (40-5300) (Invitrogen, 40-5300, 1:50); Mouse monoclonal anti-Connexin43 clone4EC2 (Millipore, MAB3067, 1:50); Rabbit monoclonal anti-α-actinin [EP2529Y] (Abcam, ab68167, 1:50); Mouse monoclonal anti-Troponin T (13-11) (Invitrogen, MA5-12960, 1:50); Rabbit monoclonal anti Claudin 5 (Acris Antibodies, DP157, 1:50); Mouse monoclonal anti-Claudin-5 (OTI1G4) (Novus Biologicals, NBP2-02259, 1:50); Rabbit polyclonal anti-SGLT2/SLC5A2 (Novus Biologicals, NBP1-92384, 1:50); Mouse anti-Caveolin-3 (BD Biosciences, 610420, 1:50); Mouse polyclonal Anti-Ryanodine Receptor [34C] (Abcam, ab2868, 1:50); Mouse monoclonal Anti-NG2 (132,38) (Santa Cruz Biotechnology, sc-33666, 1:50); Rat monoclonal Anti-Laminin-2 α [4H8-2] (Abcam, ab11576, 1:50); Mouse monoclonal Anti-β1-integrin (P4C10) (Novus Biologicals, NBP2-80818, 1:50); Mouse monoclonal anti-α-Dystroglycan (IIH6) (Santa Cruz Biotechnology, sc-53987, 1:50); Mouse monoclonal anti-NeuN clone A60 (Sigma-Aldrich, MAB377, 1:50); Mouse monoclonal anti-Calsequestrin (ThermoFisher Scientific, MA3-913, 1:50).

The secondary fluorescent antibodies used in this study, all purchased from *Invitrogen* (1:100), were as follows: Donkey anti-mouse Alexa fluor^TM^ 488 (Invitrogen, A21202), Donkey anti-rabbit Alexa fluor^TM^ 488 (Invitrogen, A21206), Goat anti-mouse Alexa fluor^TM^ 488 (Invitrogen, A11029), Donkey anti-mouse Alexa fluor^TM^ 546 (Invitrogen, A10036), Donkey anti-rabbit Alexa fluor^TM^ 568 (Invitrogen, A10042), Goat anti-rat Alexa fluor^TM^ 594 (Invitrogen, A11007) Goat anti-mouse AlexafluorTM594 (Invitrogen, A11005), Goat anti-rabbit Alexa fluor^TM^ 594 (Invitrogen, A11012), Goat anti-rat Alexa fluor^TM^ 633 (Invitrogen, A21094), Goat anti-rabbit Alexa fluor^TM^ 647 (Invitrogen, A21244).

The fluorescent probes were as follows: 4,6-diamidino-2-phenylindole (DAPI) nuclear probe (Sigma-Aldrich, 32670, 1:1000); Alexa Fluor™ 488-conjugated-wheat germ agglutinin (WGA) (ThermoFisher Scientific, W11261, 1:1000); Alexa Fluor™ 594-conjugated-WGA (ThermoFisher Scientific, W11262, 1:1000); Alexa Fluor™ 633-conjugated-WGA (ThermoFisher Scientific, W21404, 1:1000); Alexa Fluor™ 568-conjugated-Isolectin GS-IB4 (IsoB4) (ThermoFisher Scientific, I21412, 1:1000); SPY555-actin cell-permeant actin probe (Spirochrome, SC202, 1:1000); MitoTracker®Red CMXRos cell-permeant MitoTracker probe (ThermoFisher Scientific, M7512, 1:1000); Potential-Sensitive Di-8-ANEPPS dye (ThermoFisher Scientific, D3167, 1:1000).

### Immunofluorescent staining of paraffin-embedded heart sections

Formalin-fixed paraffin-embedded (FFPE) cardiac tissue sections were deparaffinized, rehydrated and subjected or not to heat-induced epitope retrieval using pH 6 Citrate buffer. The samples were then permeabilized (0.5% Triton X100-PBS), blocked (Dako Protein Block, # X0909, Dako), and stained overnight at 4°C with primary antibodies (1:100), followed by a 1-hour incubation at room temperature with fluorescent secondary antibodies in the presence of fluorescent wheat germ agglutinin (1:1000) or Isolectin GS-IB4 (IsoB4) (1:100) (all diluted in 0.1% BSA / 0.1% Tween 20 / 0.5% Triton X-100 -PBS). Nuclei were visualized with DAPI (1:2000). All images were acquired on a Zeiss LSM 900 confocal microscope using Zen blue 3.3 software (Carl Zeiss).

### Purification of adult cardiomyocytes from mouse hearts

Rreshly isolated ventricular cardiomyocytes were prepared as described previously^24^, excepted that 25 µM Blebbistatin (Clinisciences, T6038) was used for contraction inhibition instead of 2,3-Butanedione 2-monoxime (BDM)

### Western blotting

For protein analysis from cardiac extracts, cardiac tissue biopsies (1 biopsy ∼3 mg / kinetic point) or purified-adult cardiomyocytes were lysed in Lysis buffer (50 mM Tris-HCl pH 7.4, 150 mM NaCl, 1 mM EDTA, 1% TritonX100) in the presence of a protease inhibitor cocktail (Roche, Cat# 04693159001). After homogenization using a Precellys lysis kit (Bertin Instruments, #03961-1-0032 CK14), samples were centrifuged at 13,000 × g at 4°C for 10 min, and supernatants were collected. The protein concentration of extracts was determined by the Lowry method (Bio-Rad, Detergent-compatible *DC*™ Protein Assay Kit, I #5000111) and equal amounts of proteins (25, 50 µg or 100 µg) were subjected to SDS-PAGE and transferred to 0.45 µm nitrocellulose membranes (Amersham, #10600002). Blotted membranes were blocked with 5 % non-fat dry milk in PBS-0.1 % Tween-20 (blocking buffer) and hybridized with a primary antibody (anti-α actinin, Sigma-Aldrich, #A7732 or anti-GAPDH, Cell Signaling, #D16H11 or anti-ACE2, Abcam, #Ab108252 or anti-SGLT2, Novus Biologicals, #NBP1-92384) and a horse radish peroxidase (HRP)-conjugated secondary antibody (Promega, #W401B, #W402B), all diluted in the blocking buffer, for subsequent detection by chemiluminescence using the Amersham™ ECL Select™ Western Blotting Detection Reagent (GE Healthcare, RPN2235). The signal was detected using a Bio-Rad ChemiDocTM Imaging system and protein quantification was obtained by densitometric analysis using ImageQuant 5.2 software and expressed in arbitrary units (A.U.).

---

**Supplementary Figure 1.**
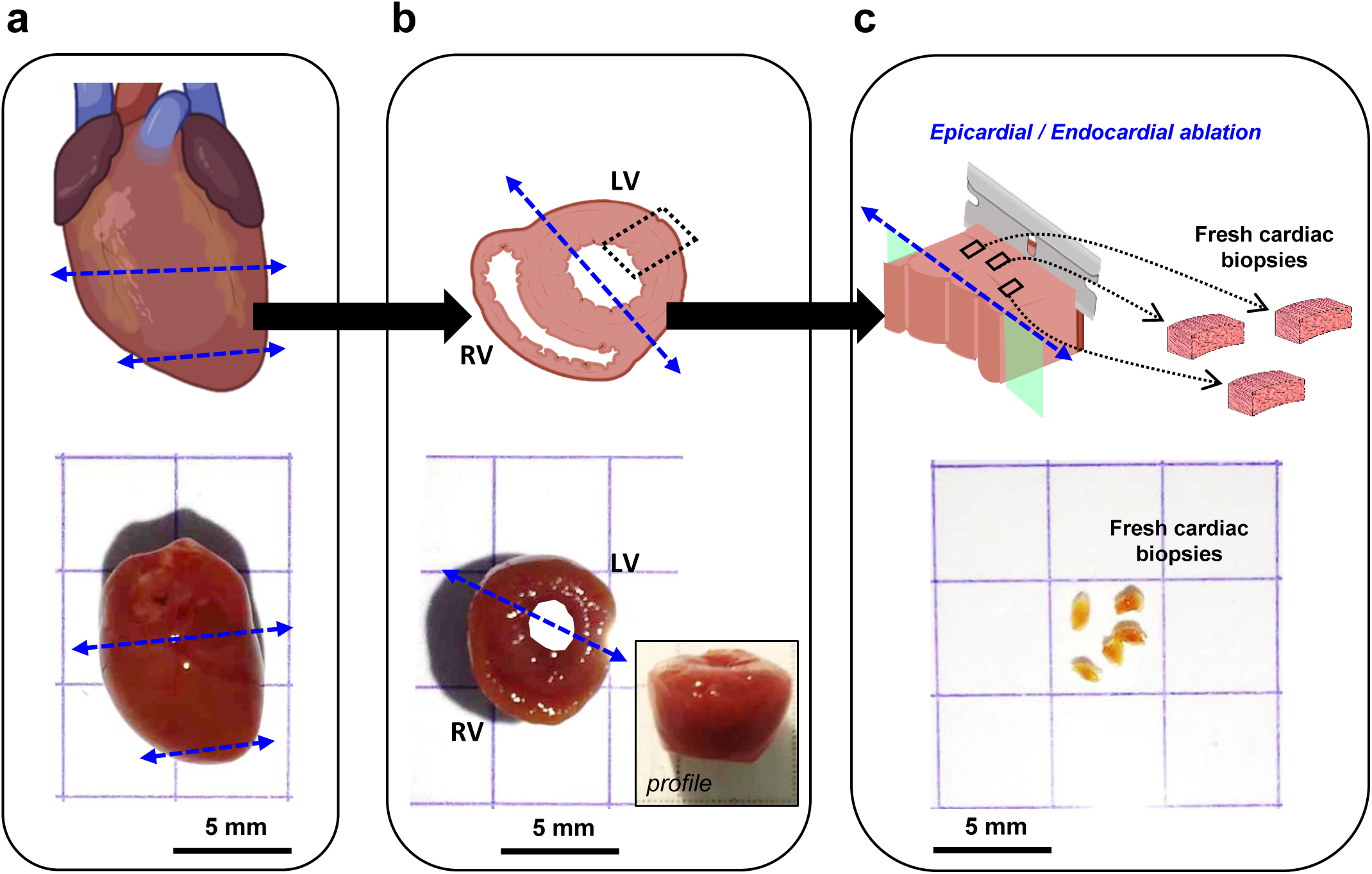
Workflow for the preparation of fresh cardiac biopsies from the left ventricle of mouse heart for the 3D-NaissI method. *Upper panels*: schematics illustrating the step-by-step process of preparation fresh cardiac biopsies from the left ventricle of the mouse heart. *Lower panels*: corresponding original photographs illustrating key stages of the procedure including (**a**) the harvested heart, (**b**) cardiac slice and (**c**) fresh biopsies. Following the extraction of the beating heart and subsequent cold PBS washing, the procedure involves: (**a)** removal of the apex and upper portion, including the atria; (**b**) obtaining a transverse slice, from which the left ventricle section is collected; (**c**) excising endocardial pillars and the epicardial layer from the left ventricular section, followed by the precise collection of 1 mm³ biopsies using a sharp, fine scalpel.

**Supplementary Figure 2.**
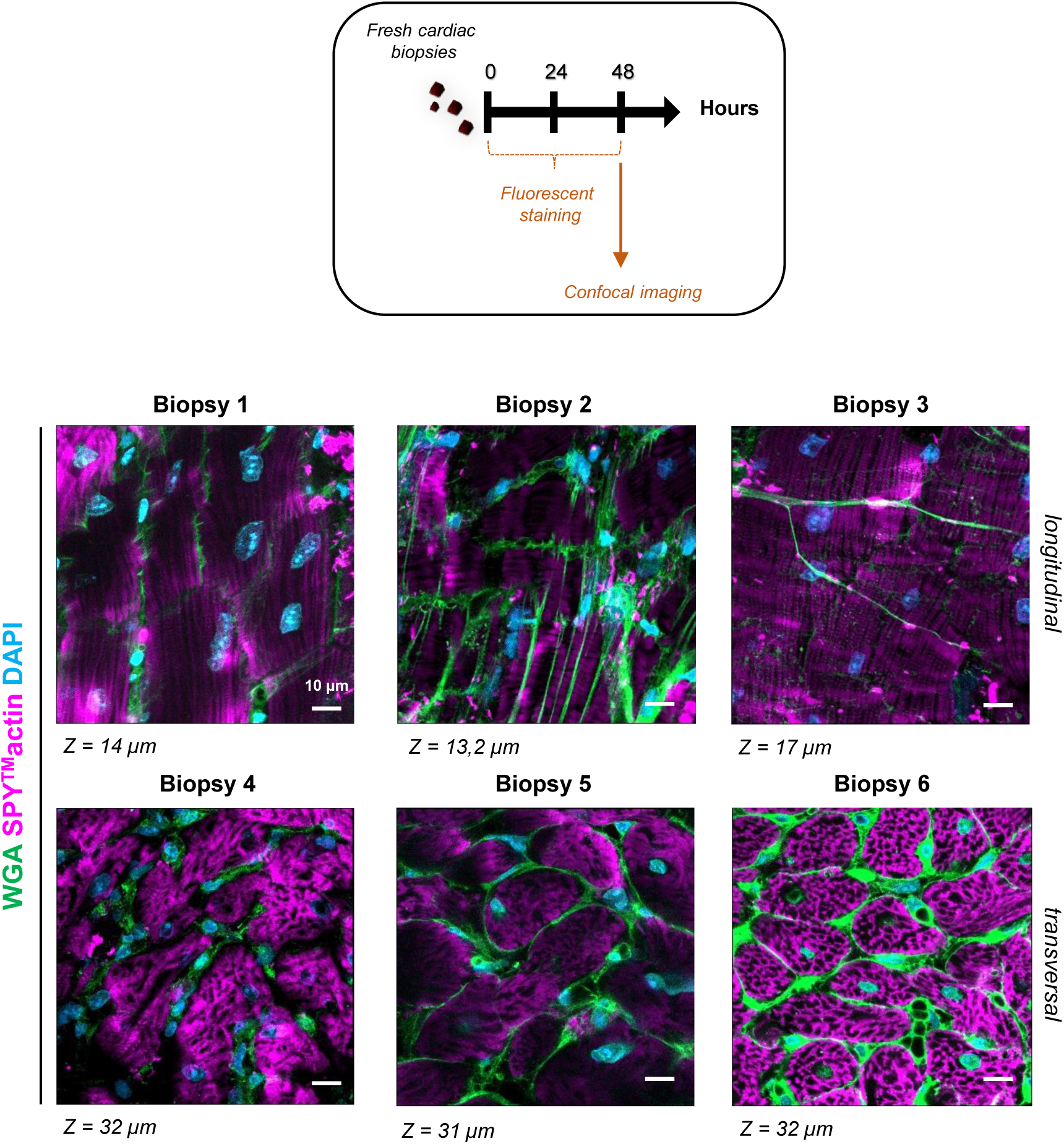
Heterogeneity of the fluorescent staining in fresh cardiac biopsies using the 3D-NaissI method. Representative confocal images of fresh cardiac biopsies from 2-month-old male mice (5 independent experiments) stained during 48 hours with fluorescent wheat germ agglutinin (WGA), SPY^TM^-actin and DAPI.

**Supplementary Figure 3.**
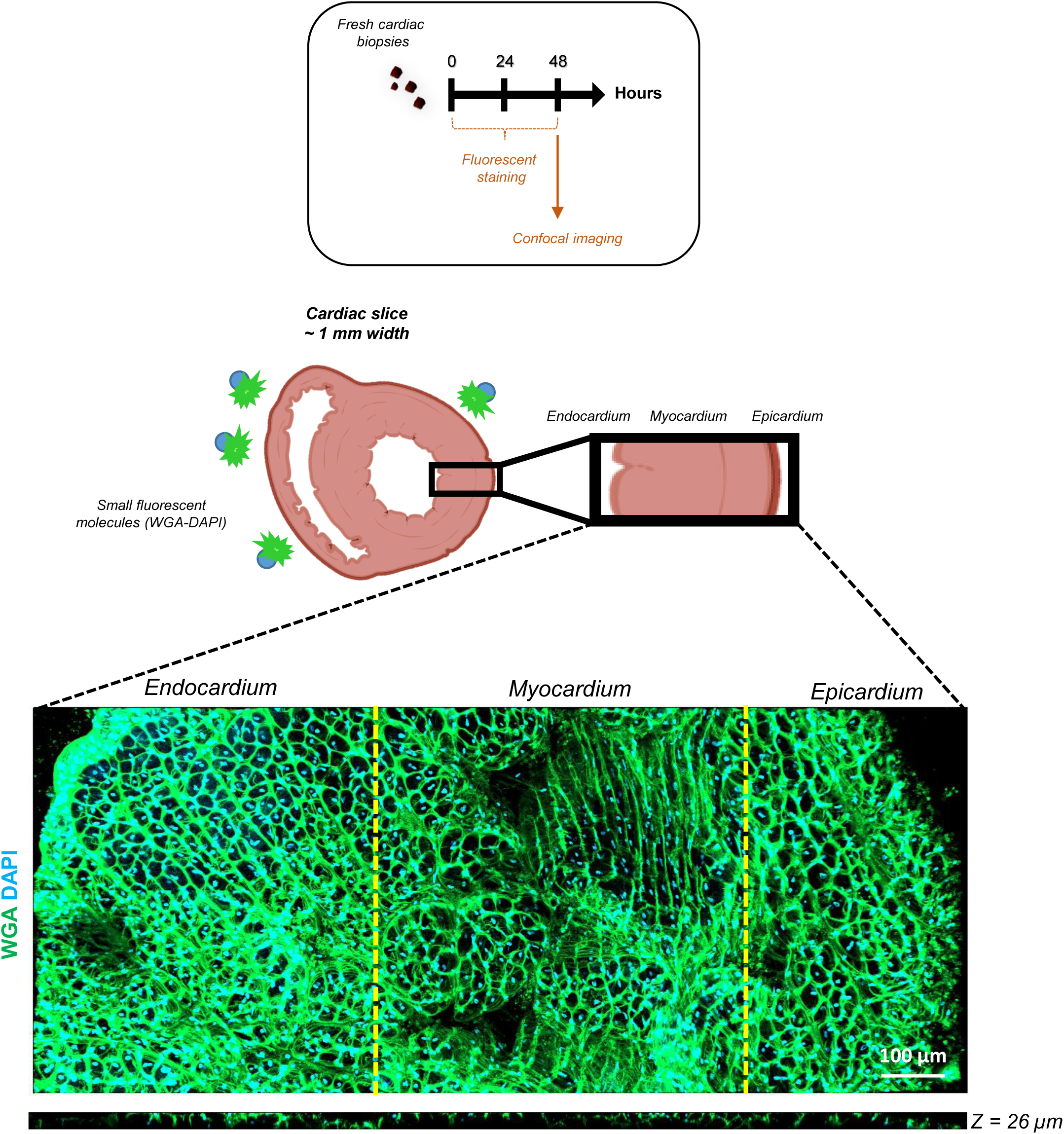
Extension of the 3D-NaissI method to fluorescent labeling of large fresh heart tissue sections. Representative 20x (air) confocal images of extensive cardiac biopsies encompassing the endocardium, myocardium and epicardium, from 2-month-old male mice (2 independent experiments) and stained for 48 hours with fluorescent wheat germ agglutinin (WGA) and DAPI.

**Supplementary Figure 4.**
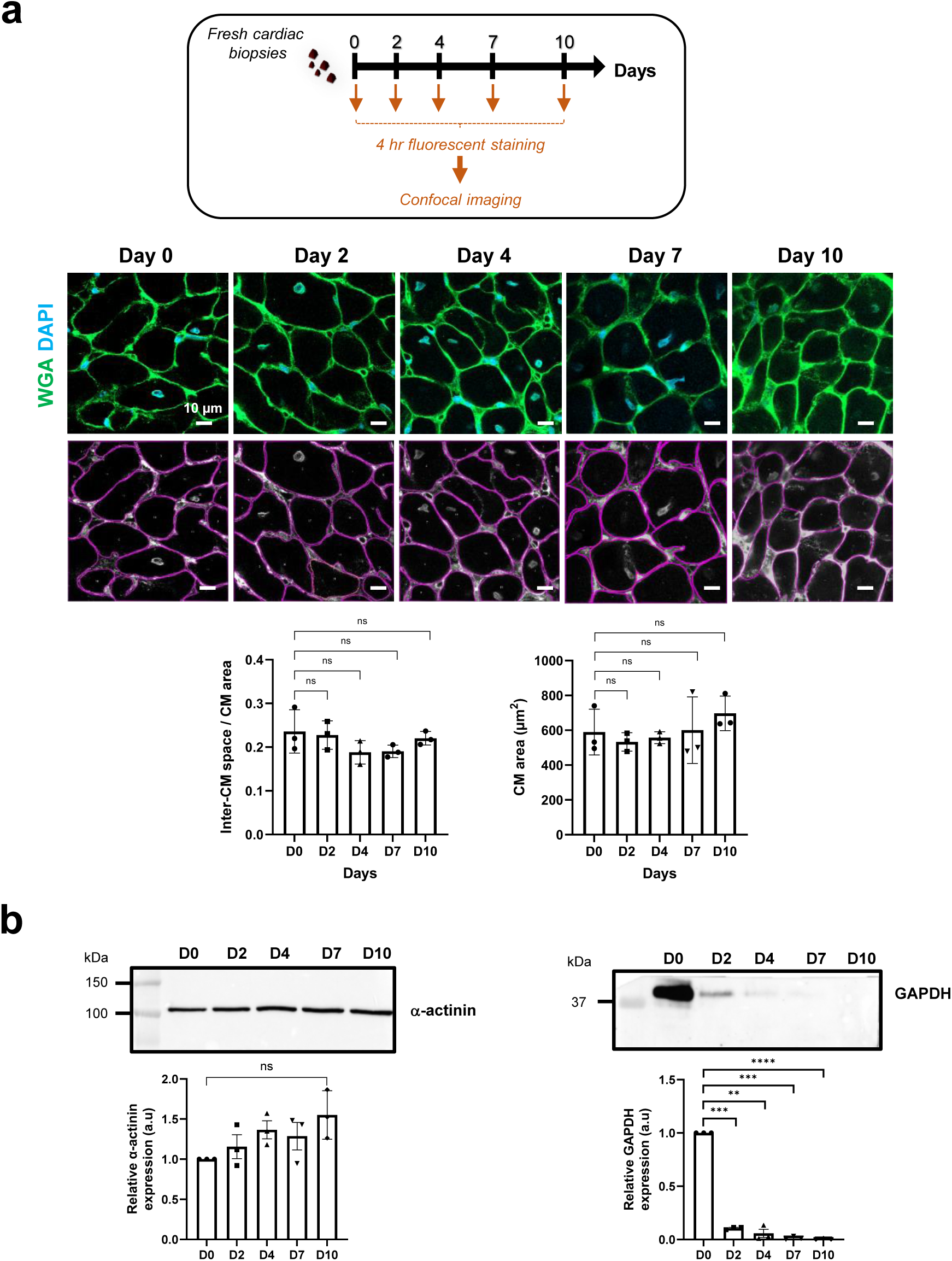
Stability kinetics of fresh cardiac biopsies. **a** Stability of fresh cardiac biopsies from 2-month-old male mice assessed at 4°C in PBS by quantification of the inter-cardiomyocyte (CM) space (indicative of the cardiac tissue cohesion) and CM area at 0, 2, 4, 7 or 10 days post-biopsy collection, following a 4 hr-fluorescent staining with wheat germ agglutinin (WGA) and DAPI before confocal imaging. Representative confocal images are presented (upper panels) with corresponding manual CM contouring using Image J (lower panels). Data are mean ± s.d.; *n*=3 mice (5-10 biopsies/mouse, 1 image /biopsy), one-way ANOVA, Dunnett’s post-hoc test with D0 as the control. **b** Quantification of α-actinin (upper panel) and GAPDH (lower panel) expression levels by western blot in lysates from fresh cardiac biopsies of 2-month-old male mice. The biopsies were preserved at 4°C in PBS for 0, 2, 4, 7 or 10 days post-collection. Data are mean ± s.e.m.; *n*=3 mice (1 biopsy ∼ 3mg /time/mouse, 1 image /biopsy), one-way ANOVA, Dunnett’s post-hoc test with D0 as the control.

**Supplementary Figure 5.**
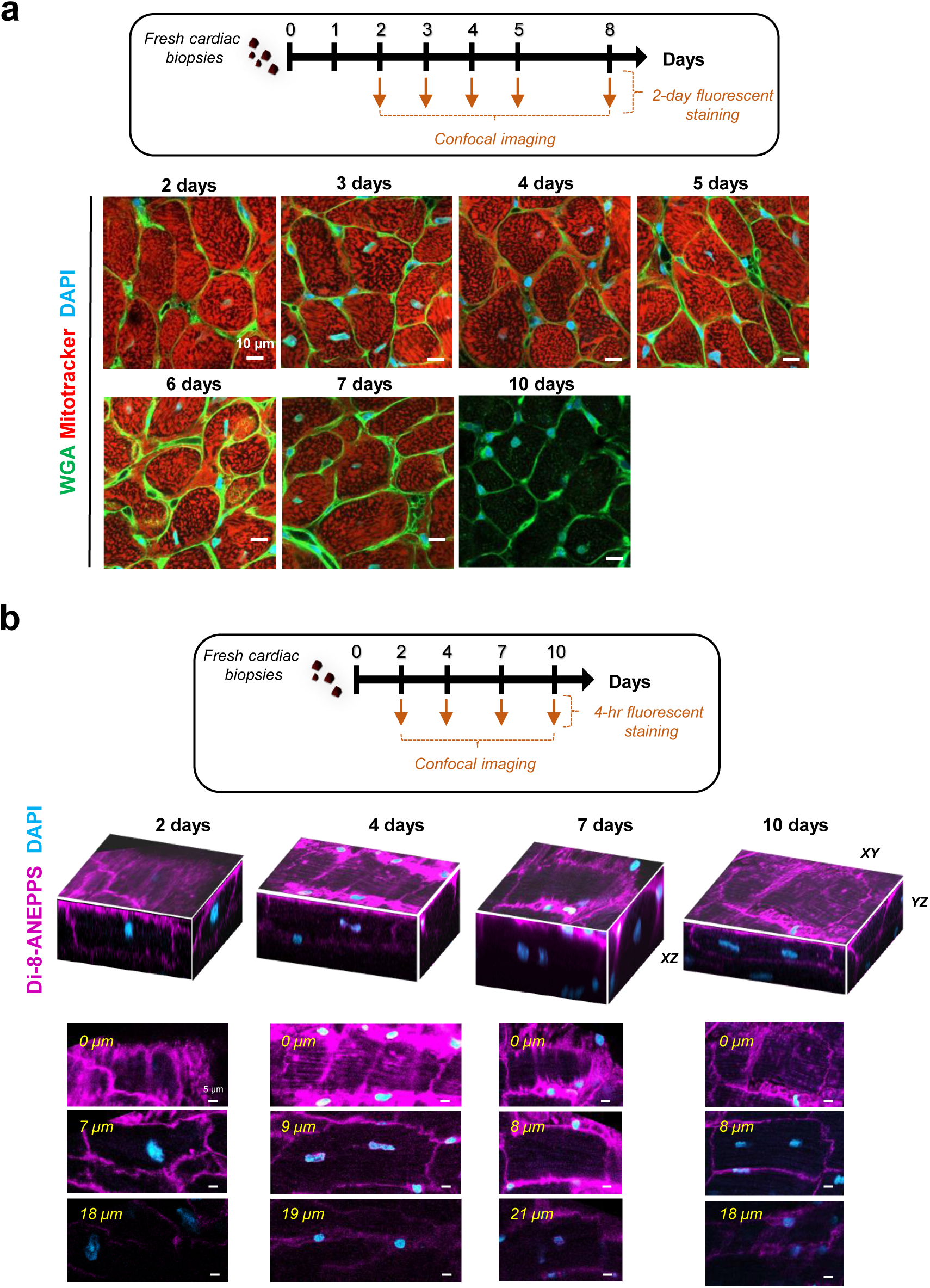
Viability kinetics of fresh cardiac biopsies. **a** Representative confocal images (3 independent experiments; 5-6 biopsies/time) depicting the viability of cardiomyocyte mitochondria, in fresh cardiac biopsies from 2-month-old male mice, and assessed through a 2-day co-staining with fluorescent wheat germ agglutinin (WGA)-Mitotracker and DAPI after preservation at 4°C in PBS for 2-10 days post-collection. **b** Representative confocal images (2 independent experiments; 5-6 biopsies/time) showing XY, XZ, YZ projections, and associated Z-stacks of Di-8-ANEPPS-fluorescent-positive cardiomyocytes (whole-living cardiomyocytes) in fresh cardiac biopsies from 2-month-old male mice. The biopsies were preserved at 4°C in PBS for 2-10 days post-collection following a 4-hour Di-ANEPPS staining before confocal imaging.

**Supplementary Figure 6.**
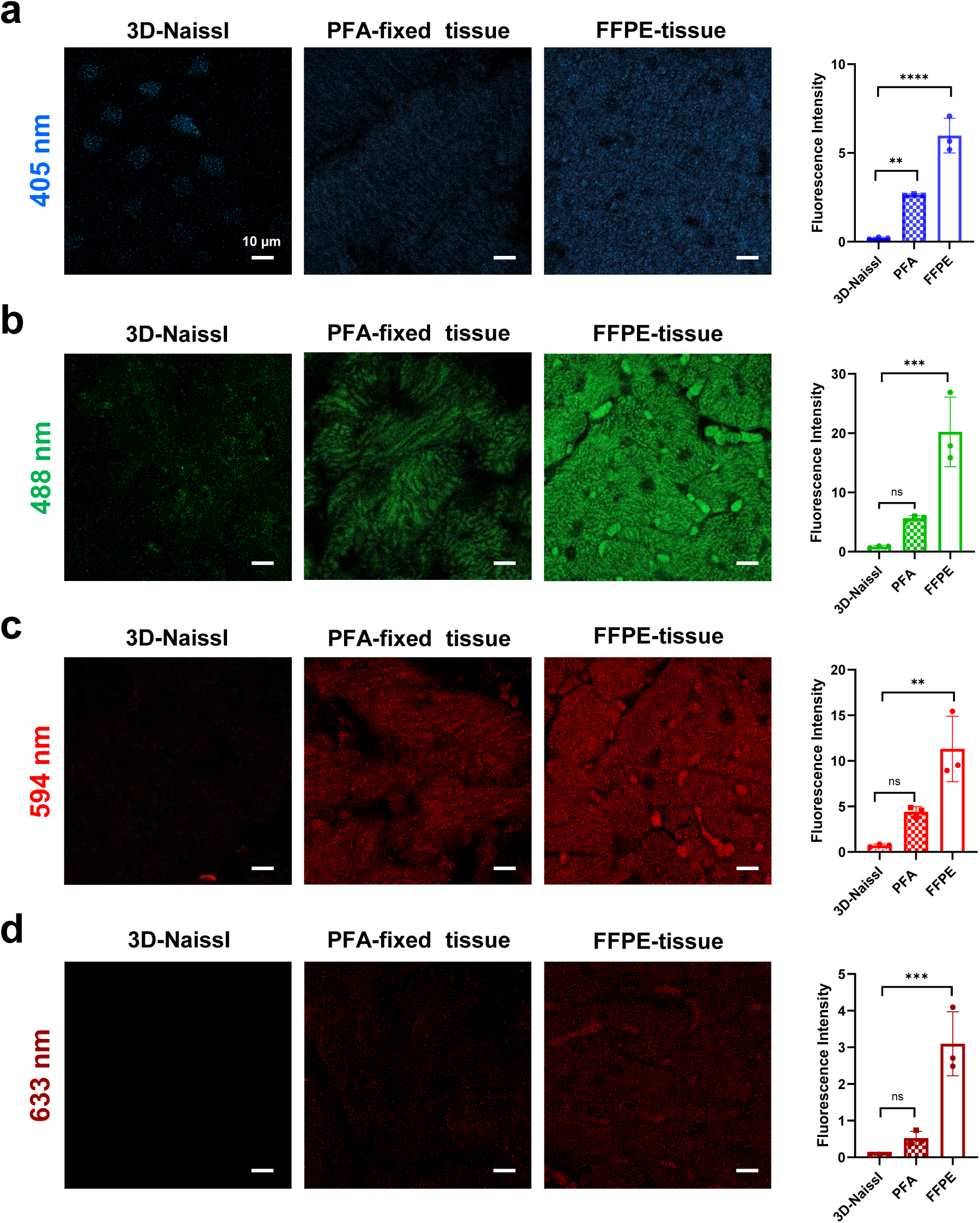
Performance of 3D-NaissI tissue autofluorescence compared to conventional histological techniques. Representative confocal imaging (left panels) and corresponding quantification of autofluorescence (right panels) (3 independent experiments) in fresh (3D-NAissI), paraformaldehyde (PFA)-fixed or Formalin-Fixed Paraffin-Embedded (FFPE) cardiac tissue obtained from 2-month-old male mice. Autofluorescence signals were captured at (**a**) 353 nm excitation and 380-450 nm registration (“blue” channel), (**b**) at 480 nm excitation and 460-510 nm registration (“green” channel), (**c**) at 590 nm excitation and 510-620 nm registration (“red” channel) or (**d**), at 631 nm excitation and 510-620 nm registration (“far red” channel). Data are mean ± s.e.m.; *n*=3 mice, (5-8 biopsies/mouse, 1 image /biopsy); one-way ANOVA, Dunnett’s post-hoc test using 3D-NAissI as the control.

**Supplementary Figure 7.**
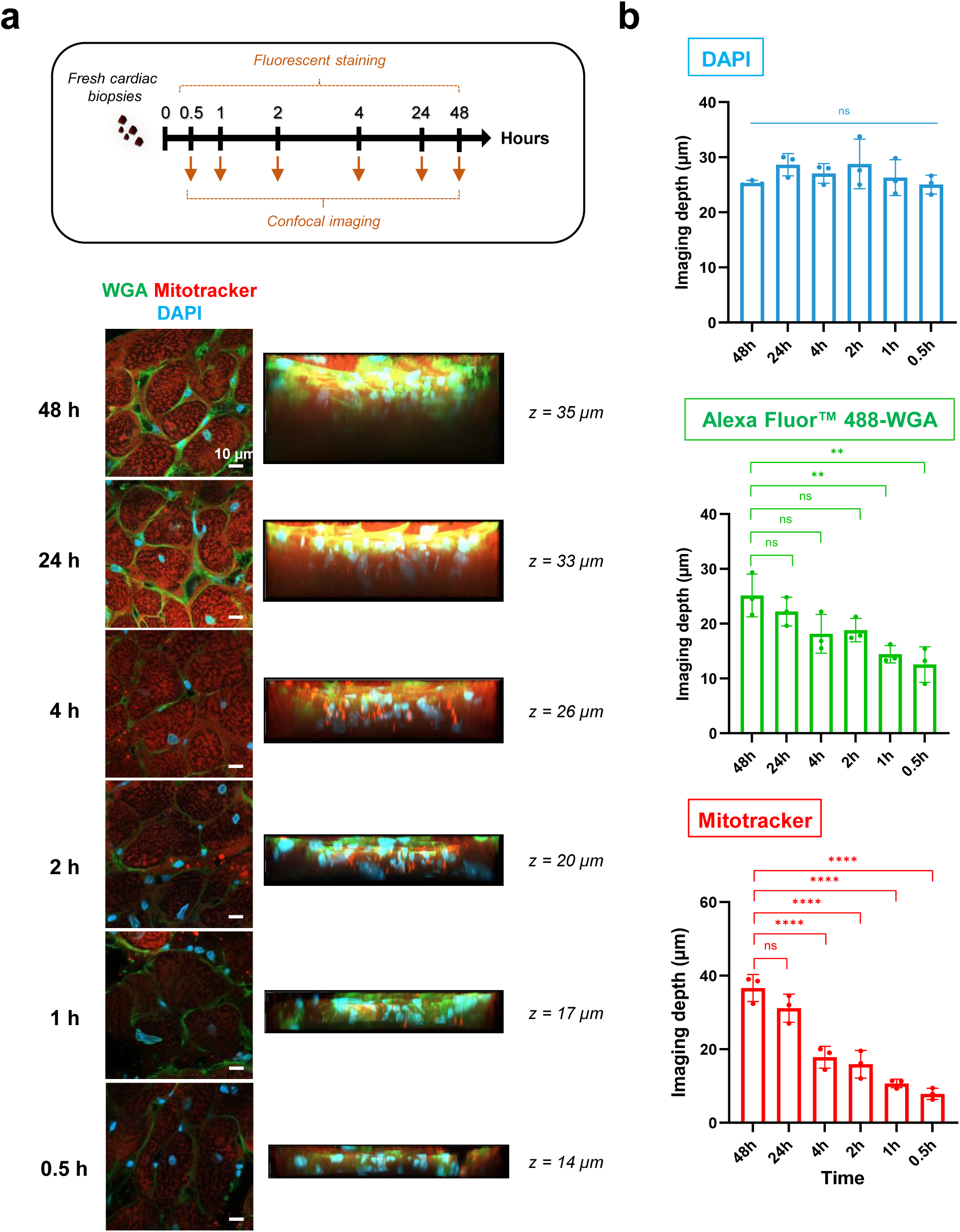
Optimization of fluorescent-small molecule staining of fresh cardiac biopsies. (**a**) Representative confocal imaging depth and (**b**) corresponding quantifications of DAPI, Alexa Fluor™ 488-conjugated-wheat germ agglutinin (WGA) and MitoTracker®Red CMXRos staining kinetics on fresh cardiac biopsies from 2-month-old male mice ranging from 0.5 to 48 hours post-biopsy collection before confocal imaging. Data are mean ± s.d. *n*=3 mice (5-10 biopsies/mouse, 1 image /biopsy), one-way ANOVA, Dunnett’s post-hoc test using 0.5 hour as the control.

**Supplementary Figure 8.**
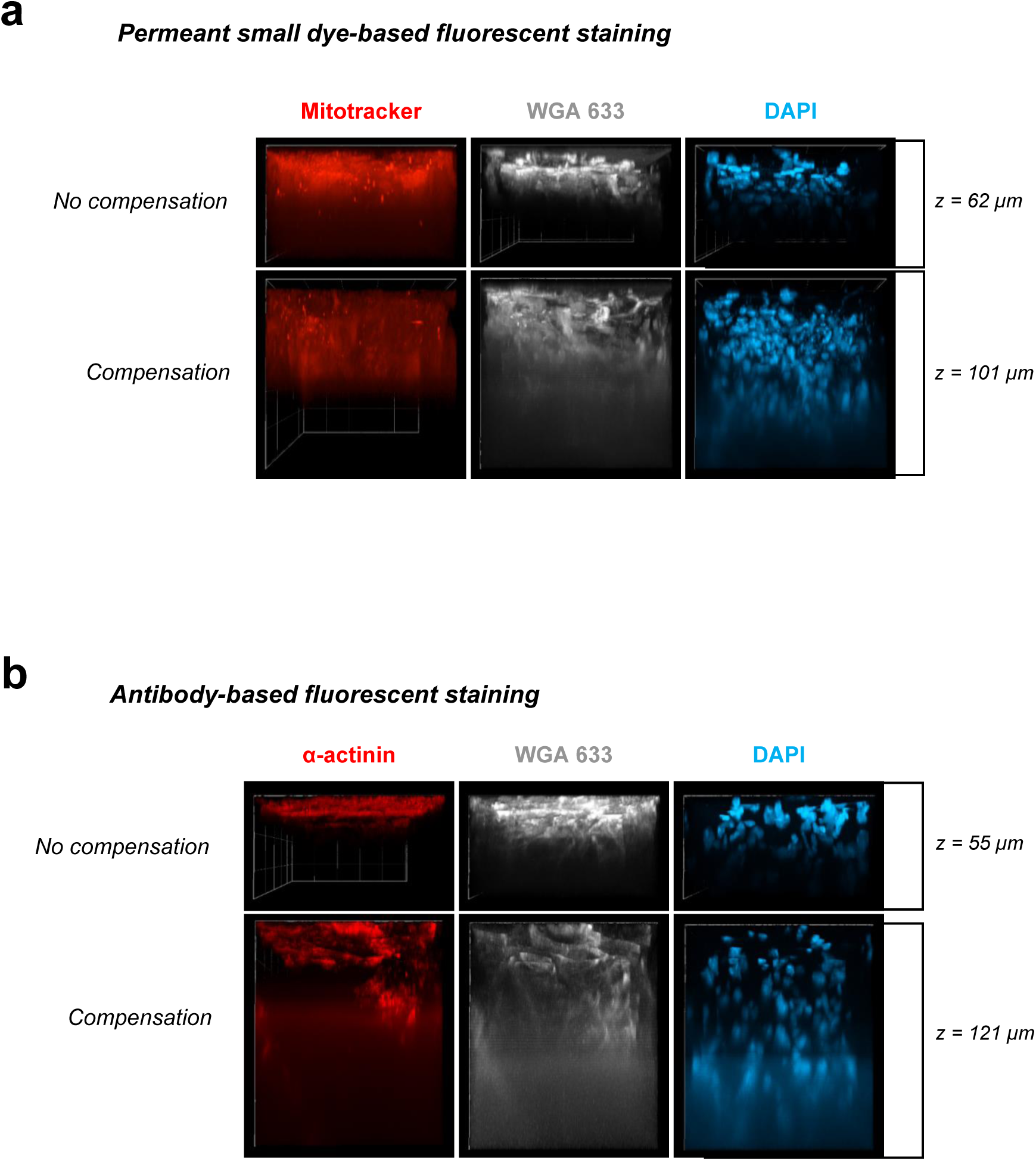
Optimization of fluorescent staining depth in fresh cardiac biopsies. Representative confocal imaging depth (2 independent experiments) of fresh cardiac biopsies from 2-month-old male mice stained with (**a**) small fluorescent molecules MitoTracker®Red CMXRos, Alexa Fluor™ 633-conjugated-wheat germ agglutinin (WGA) and DAPI, or (**b**) anti-α-actinin antibody/anti Goat anti-rabbit Alexafluor^TM^594 and Alexa Fluor™ 633-WGA, and showing in both cases image acquisition with (lower panels) or without (upper panels) laser compensation (Laser power increase every 10 µm).

**Supplementary Figure 9.**
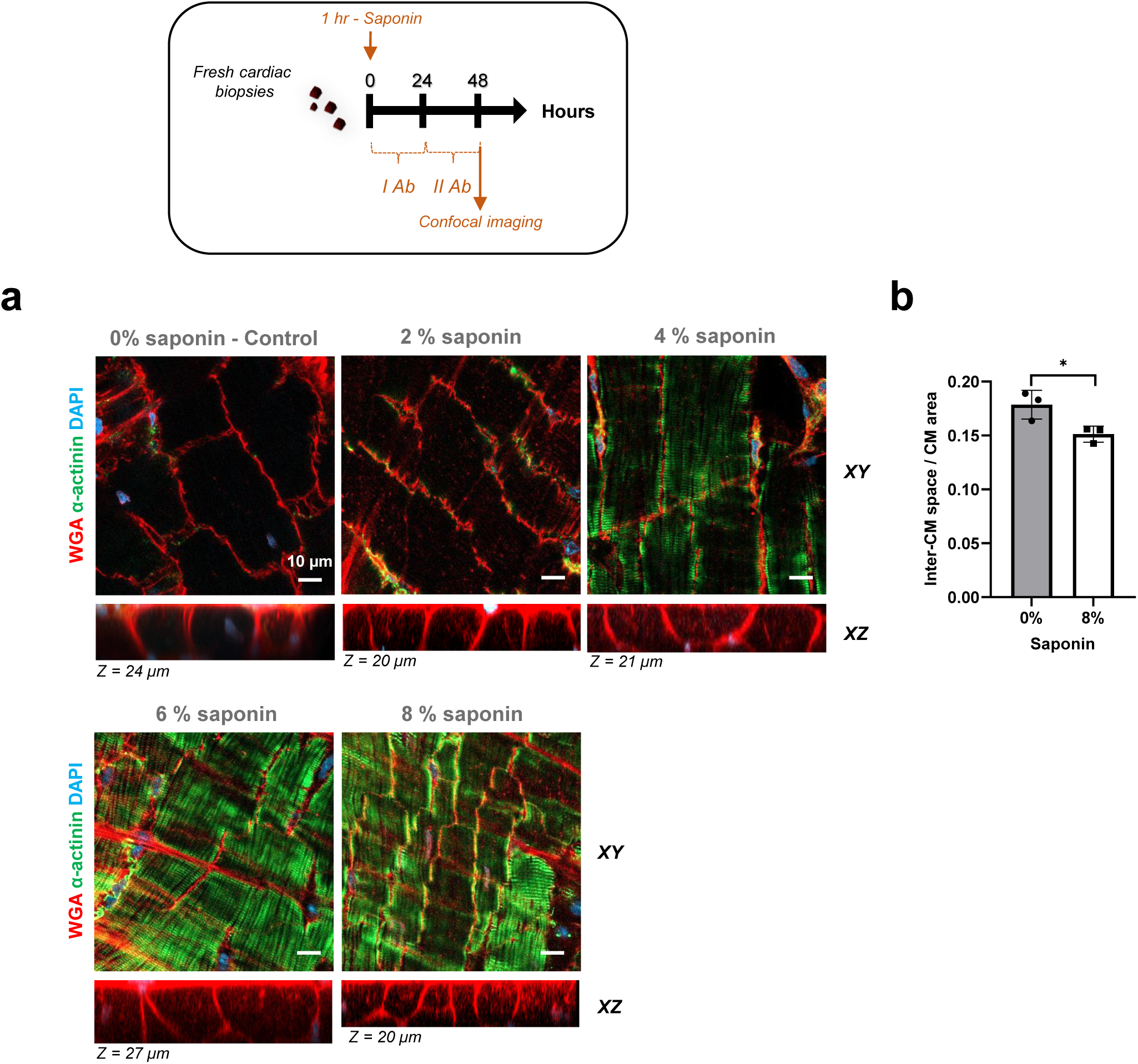
Optimization of antibody-cell permeability for fluorescent staining of fresh cardiac biopsies. **(a)** Representative confocal images (3 independent experiments) and (**b)** corresponding quantification of the inter-cardiomyocyte (CM) space (indicative of the cardiac tissue cohesion), in fresh cardiac biopsies from 2-month-old male mice. Biopsies were treated or not (0% saponin-Control) with different saponin concentrations for 1 hour at 4°C in PBS, followed by 24 hours of successive incubation with anti-α-actinin antibody, conjugated-wheat germ agglutinin (WGA), DAPI, and an additional 24 hours with the II fluorescent antibody before confocal imaging. XZ projections indicate the imaging of entire cardiomyocytes. Data are mean ± s.d. n=3 mice (5-10 biopsies/mouse, 1 image /biopsy), unpaired Student’s t-test.

**Supplementary Figure 10.**
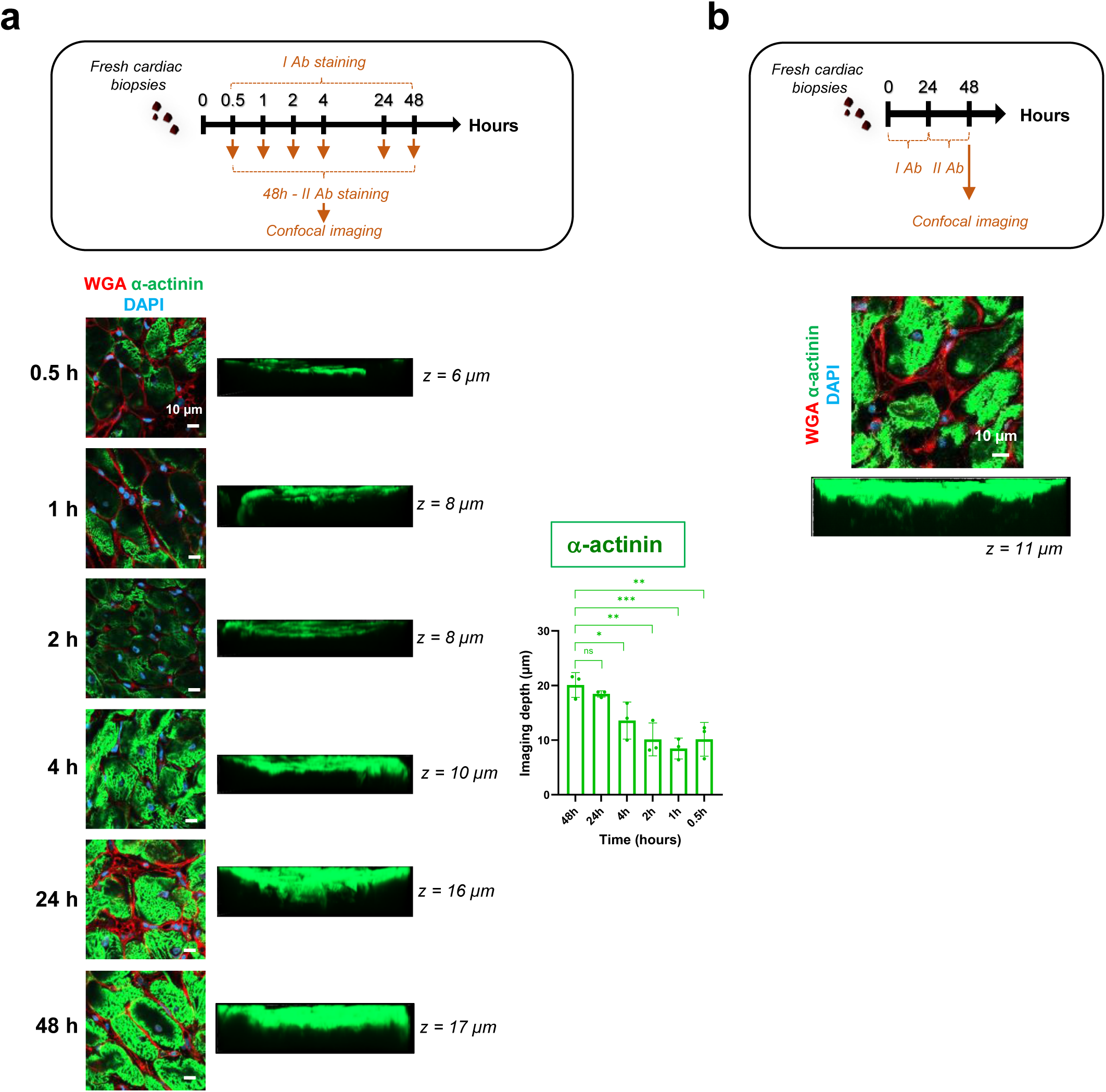
Optimization of fluorescent-antibody staining in fresh cardiac biopsies. **a** Representative confocal imaging depth (left panels) and corresponding quantifications (right panels) of DAPI, Alexa Fluor™ 594-conjugated-wheat germ agglutinin (WGA) and anti-α-actinin staining kinetics on fresh cardiac biopsies from 2-month-old male mice. Staining periods ranged from 0.5 to 48 hours, followed by a 48-hour staining with the secondary fluorescent against anti-α-actinin antibody before imaging. Data are mean ± s.d. *n*=3 mice (5-10 biopsies/mouse, 1 image /biopsy), one-way ANOVA, Dunnett’s post-hoc test with 48 hours as the control. **b** Similar experiment to **(a)**, with a reduced incubation time to 24 hours with the primary antibody anti-α-actinin and 24 hours with the secondary complementary fluorescent antibody. A representative image is shown.

**Supplementary Figure 11.**
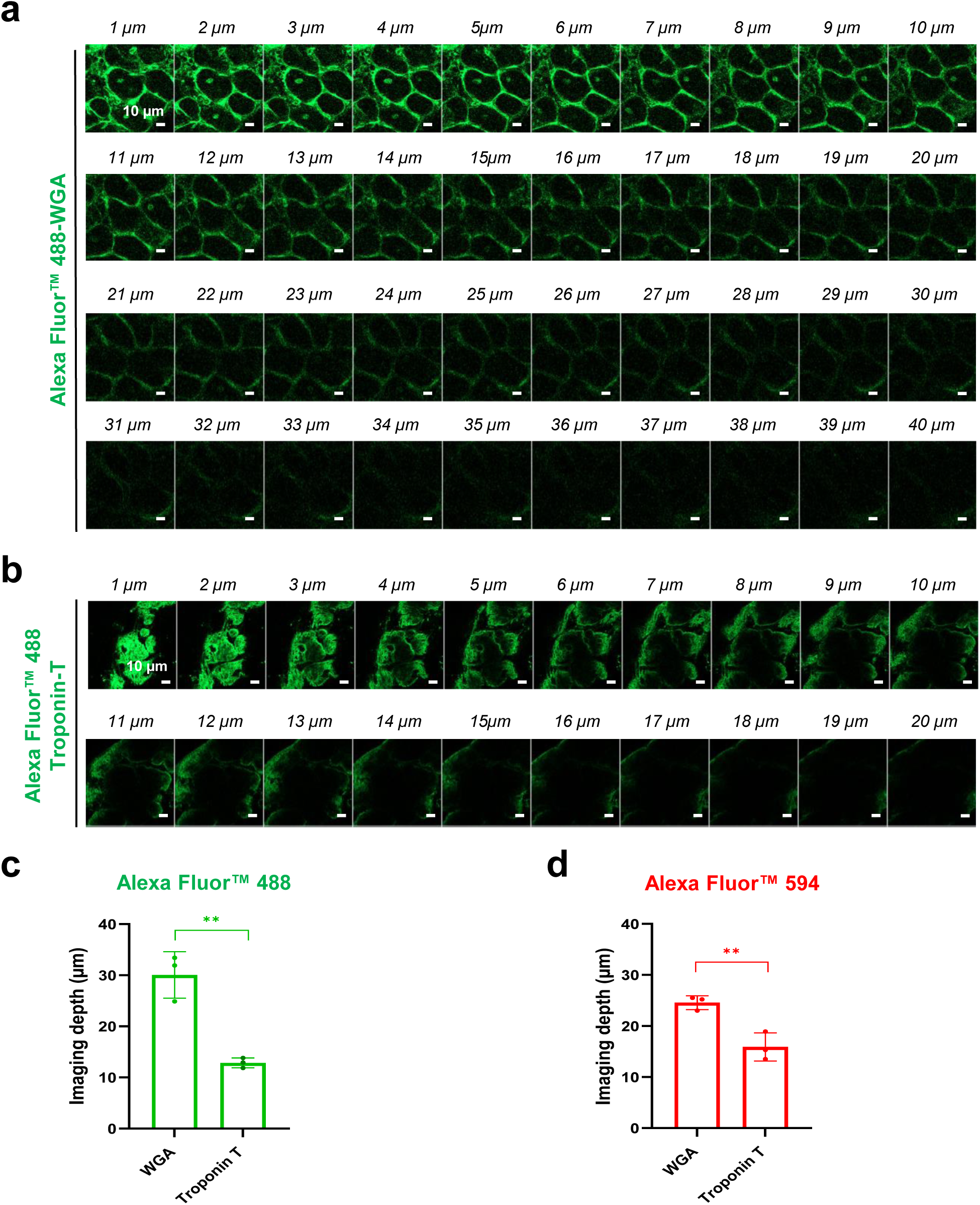
Fluorescent labeling depth independence on laser excitation wavelength, determined by fluorescent molecule size. **a, b, c** Representative images (**a, b**) and corresponding quantifications (**c**) of imaging depth after a 48-hour staining of fresh cardiac biopsies from 2-month-old male mice with Alexa Fluor™ 488-conjugated-wheat germ agglutinin (WGA) (**a**) or anti-Troponin T/Alexa Fluor™ 488-secondary antibody (**b**). **d** quantification of imaging depth following 48-hour staining of fresh cardiac biopsies from 2-month-old male mice with Alexa Fluor™ 594-conjugated-wheat germ agglutinin (WGA) or anti-Troponin T/Alexa Fluor™ 594-secondary antibody. Data are mean ± s.d. *n*=3 mice (5 biopsies/mouse, 1 image /biopsy), unpaired Student’s t-test.

**Supplementary Figure 12.**
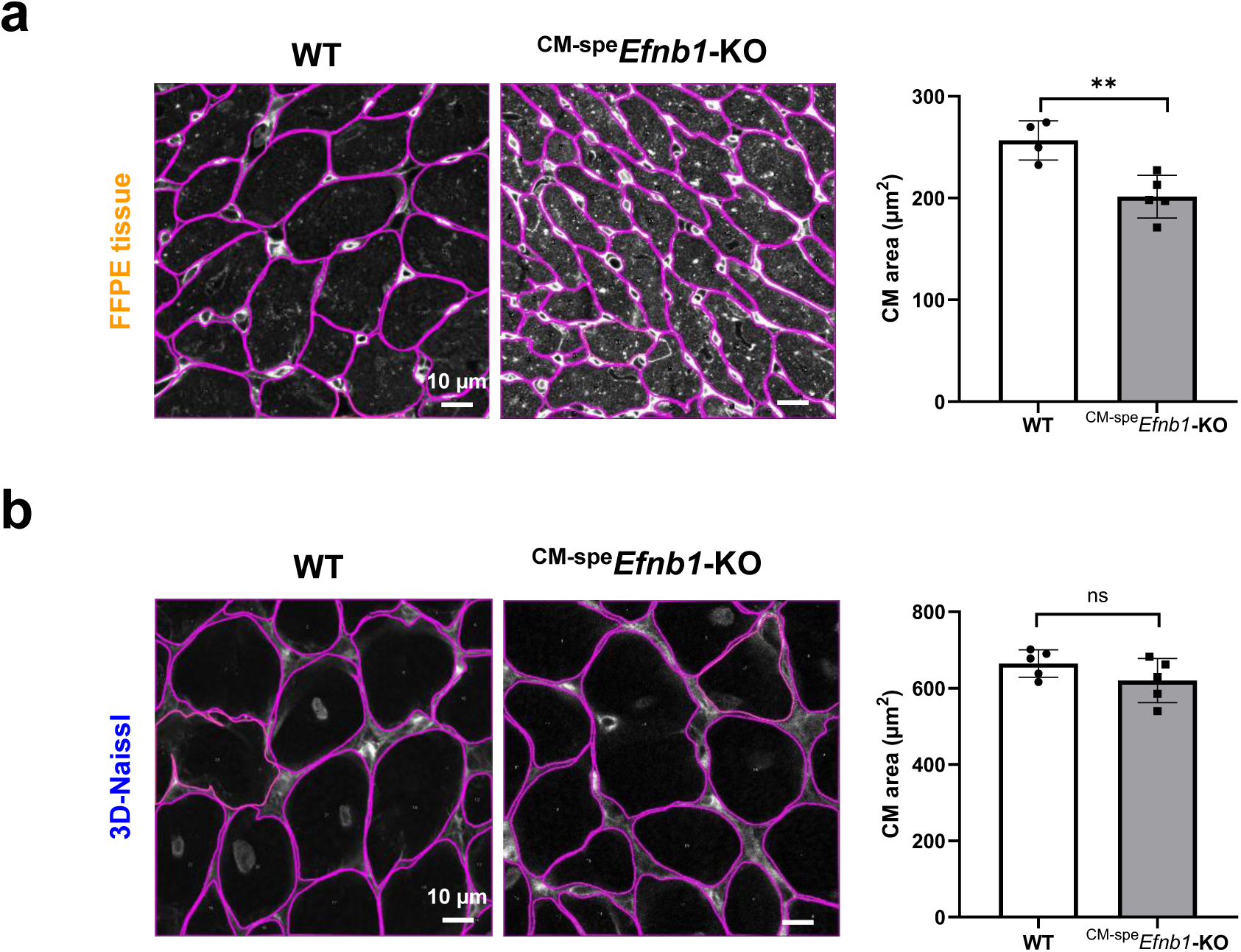
Preservation of native cardiomyocyte size by the 3D-NaissI method in different cardiac tissue phenotypes compared to conventional histological methods. **a, b** Representative manual contouring of cardiomyocyte (CM) surfaces (left panels) and corresponding CM area quantifications (right panels) performed on confocal images of fluorescent wheat germ agglutinin (WGA)-stained cardiac tissue from 2-month-old male WT or ^CM-spe^*Efnb1*-KO mice using (**a**) Formalin-Fixed Paraffin-Embedded (FFPE) cardiac tissue or **(b)** fresh cardiac biopsies (3D-NaissI method). Data are mean ± s.d. *n*=4-5 mice (8-11 biopsies/mouse, 1 image /biopsy), unpaired Student’s t-test.

**Supplementary Figure 13.**
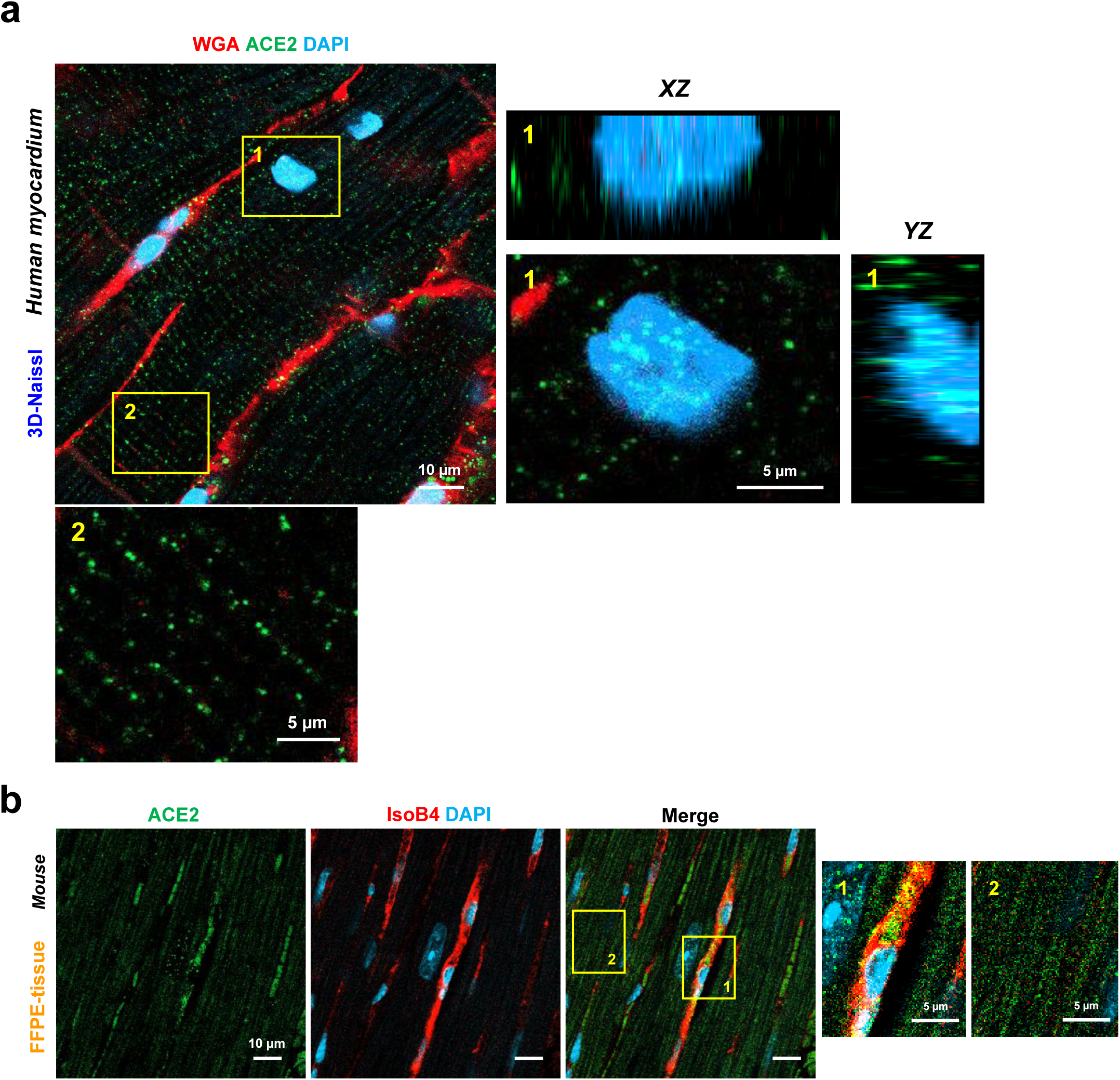
Comparison of ACE2 cardiac expression using 3D-NAissI or conventional histological method. Representative confocal imaging of: **a** ACE2 immunostaining with fluorescent wheat germ agglutinin (WGA) and DAPI in fresh cardiac tissue (3D-NaissI) from a 19-year-old man. Zoomed-in images highlight ACE2 localization within cardiomyocytes (CM) (2) and CM nuclei (1). **b** ACE2-Isolectin B4 (IsoB4)-DAPI co-staining (3 independent experiments) in Formalin-Fixed Paraffin-Embedded (FFPE) tissue from 2-month old male mice. Zoomed-in images highlight specific ACE2 localization within vascular cells (Iso-B4 positive cells) (1) but not in CMs (2).

**Supplementary Figure 14.**
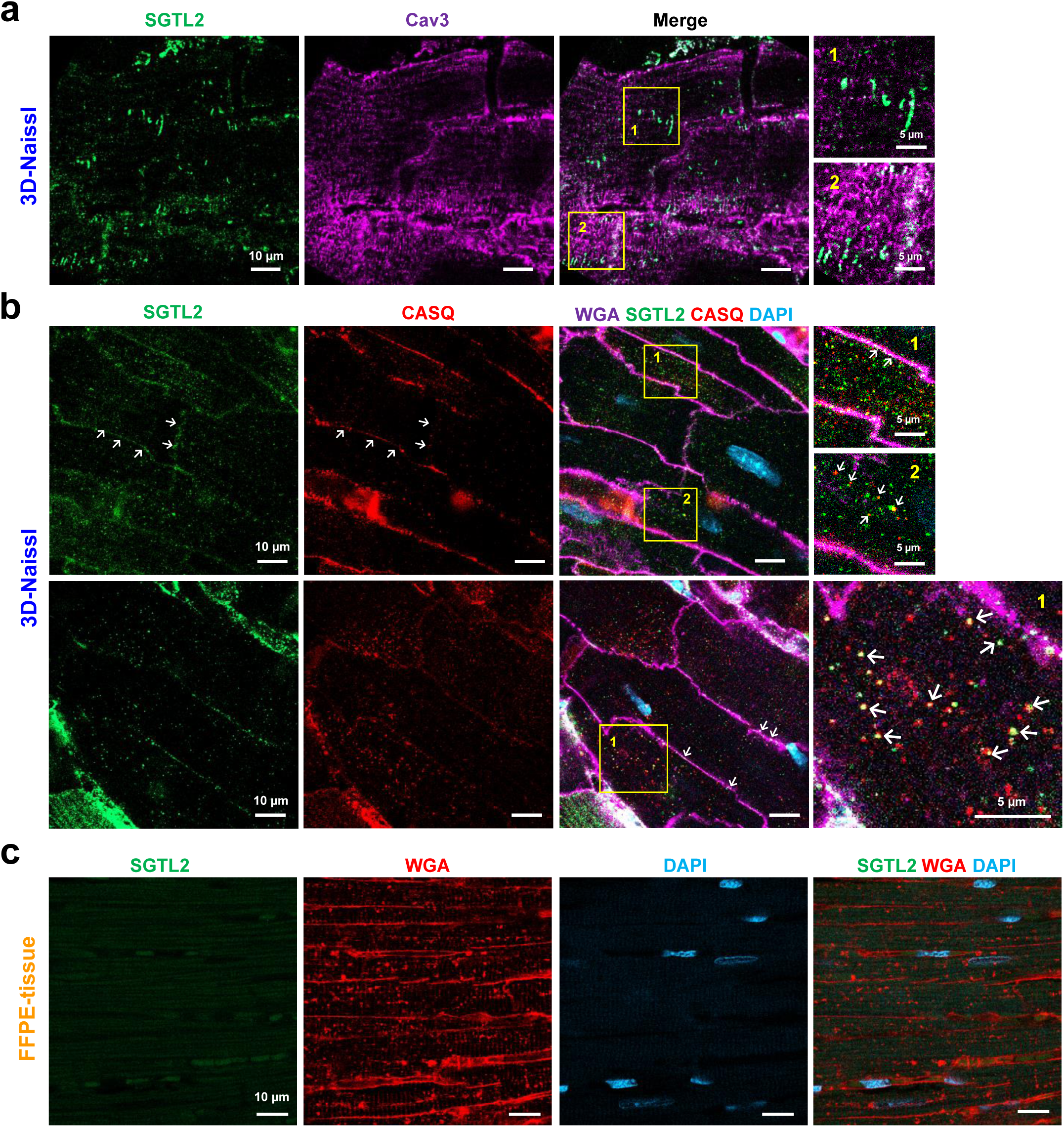
Comparison of SGTL2 cardiac expression using 3D-NAissI and conventional histological methods. Representative confocal imaging (3 independent experiments) of: **a,** SGTL2-Caveolin 3 (Cav 3)-DAPI or **b,** SGTL2-fluorescent wheat germ agglutinin (WGA)-Calsequestrin (CASQ)-DAPI or **c**, SGTL2-WGA co-stainings in fresh (3D-NAissI) (**a**, **b**) or (**c**) Formalin-Fixed Paraffin-Embedded (FFPE) tissue from 2-month old male mice. Zoomed-in images highlight specific SGTL2 staining within cardiomyocytes (CMs) (**a**), co-localization with Calsequestrin sarcoplasmic reticulum protein (**b**) but not with caveolin-3 (**a**). No specific SGTL2 staining in FFPE-cardiac tissue (**c**).

**Supplementary Figure 15.**
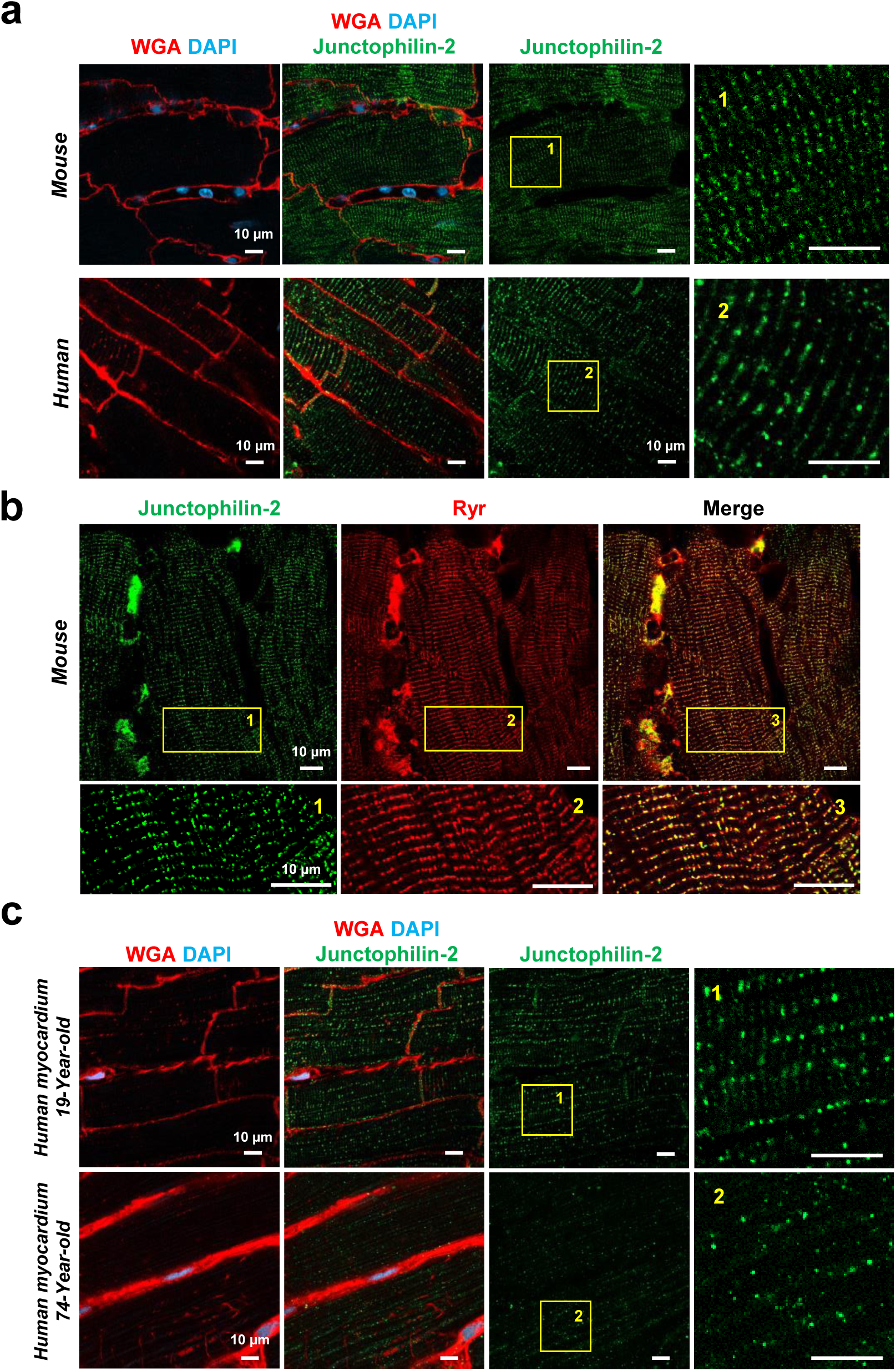
3D-NaissI visualization of key components of cardiomyocyte Excitation-Contraction (E-C) coupling. **a** Representative confocal imaging (3 independent experiments) of Junctophilin-2 co-staining with fluorescent wheat germ agglutinin (WGA) and DAPI in fresh cardiac biopsies from 2-month old male mice (upper panels) or from a 19-year-old man (lower panels). **b** Representative high resolution confocal imaging of Junctophilin-2 co-staining with Ryanodin receptor (Ryr) in fresh cardiac biopsies from 2-month old male mice, acquired using Airyscan technology. Zoomed-in images highlight the detailed spatial organization and co-localization of Junctophilin-2 and Ryr within cardiomyocytes. **c** Representative confocal imaging of Junctophilin-2 co-staining with fluorescent WGA and DAPI in fresh cardiac biopsies from a 19-year-old and a 74-year-old man. Zoomed-in images highlight differences in Junctophilin-2 spatial organization within cardiomyocytes in young versus elderly individuals.

**Supplementary Figure 16.**
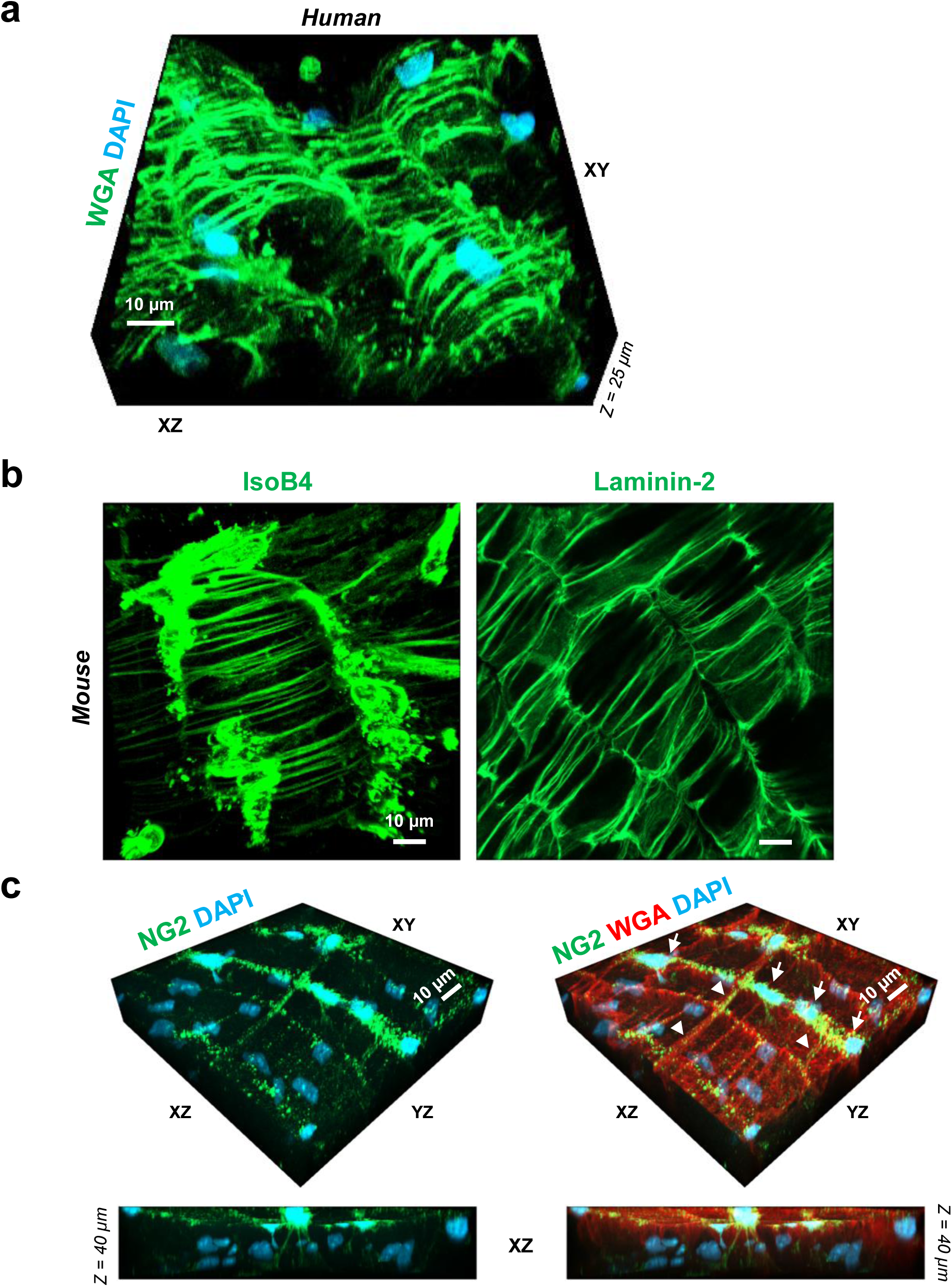
3D-NaissI visualization of a novel ring-like cardiac structure. Representative confocal 3D imaging of: **a** Fluorescent wheat germ agglutinin (WGA) AND DAPI staining using 3D-NAissI in fresh cardiac biopsies from a 26-year-old man, revealing a periodic ring-like structure along cardiac myofibrils. **b** Fluorescent Isolectin B4 (IsoB4) or laminin-2 positive-staining of the ring-like structures in fresh cardiac biopsies from 2-month old male mice. **c** Neuron-glial antigen 2 (NG2 proteoglycan) co-staining with fluorescent WGA and DAPI in fresh cardiac biopsies from 2-month old male mice, showing NG2-positive staining of ring-like structures and emanating from cell extensions (nuclei) positioned on either side of the ring-like structures along the myofibrils. XZ projection emphasizes NG2-positive cells and their cytoplasmic extensions.

**Supplementary Figure 17.**
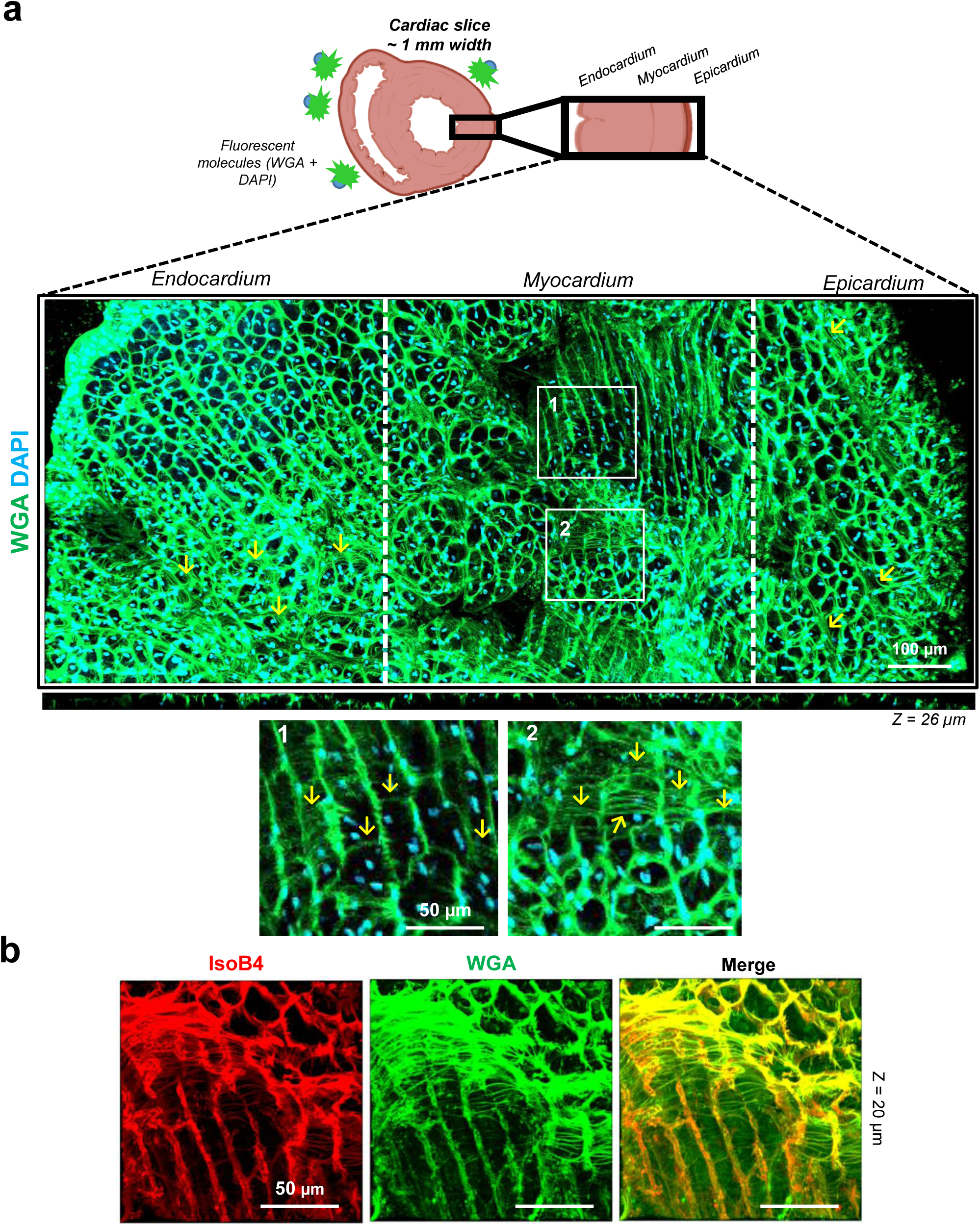
3D-NaissI visualization of ring-like cardiac structures across all left ventricle sub-compartments. Representative 20x (air) confocal images of extensive cardiac biopsies covering the left ventricle endocardium, myocardium and epicardium from 2-month-old male mice (2 independent experiments), and stained for 48 hours with fluorescent wheat germ agglutinin (WGA) and DAPI (**a**) or Isolectin-B4 (IsoB4) and WGA (**b**). Zoomed-in images highlight the presence of ring-like structures surrounding myofibrils within all three left ventricle sub-compartments (yellow arrows), exhibiting complete co-localization with IsoB4.

**Supplementary Figure 18.**
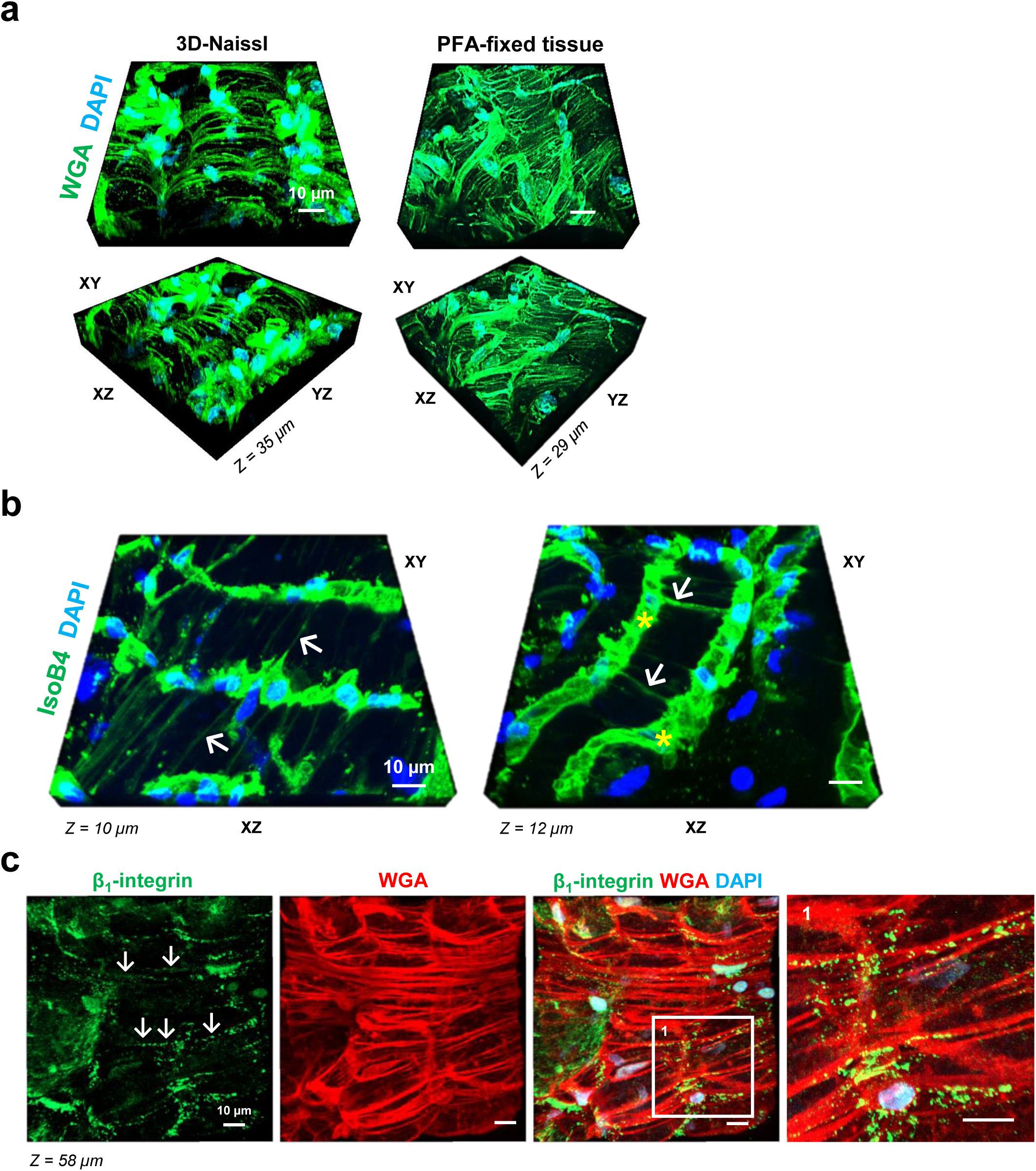
Insight into 3D Ring-like cardiac structures with the 3D-NaissI method. Representative confocal 3D imaging (3 independent experiments) of the cardiac tissue from 2-month-old male mice following: **(a)** fluorescent wheat germ agglutinin (WGA) and DAPI staining in fresh (3D-NaissI) or PFA-fixed cardiac tissue or (**b)**, fluorescent Isolectin-B4 (IsoB4) and DAPI staining in fresh cardiac biopsies. **b**, white arrows indicate the ring-like structures emanating from cells lying on lateral capillaries parallel with cardiomyocytes, **c** β_1_-integrin, WGA, DAPI co-staining in fresh cardiac biopsies showing meticulously organized β_1_-integrin-postive staining along ring-like structures at the cardiomyocyte surface.

**Supplementary Figure 19.**
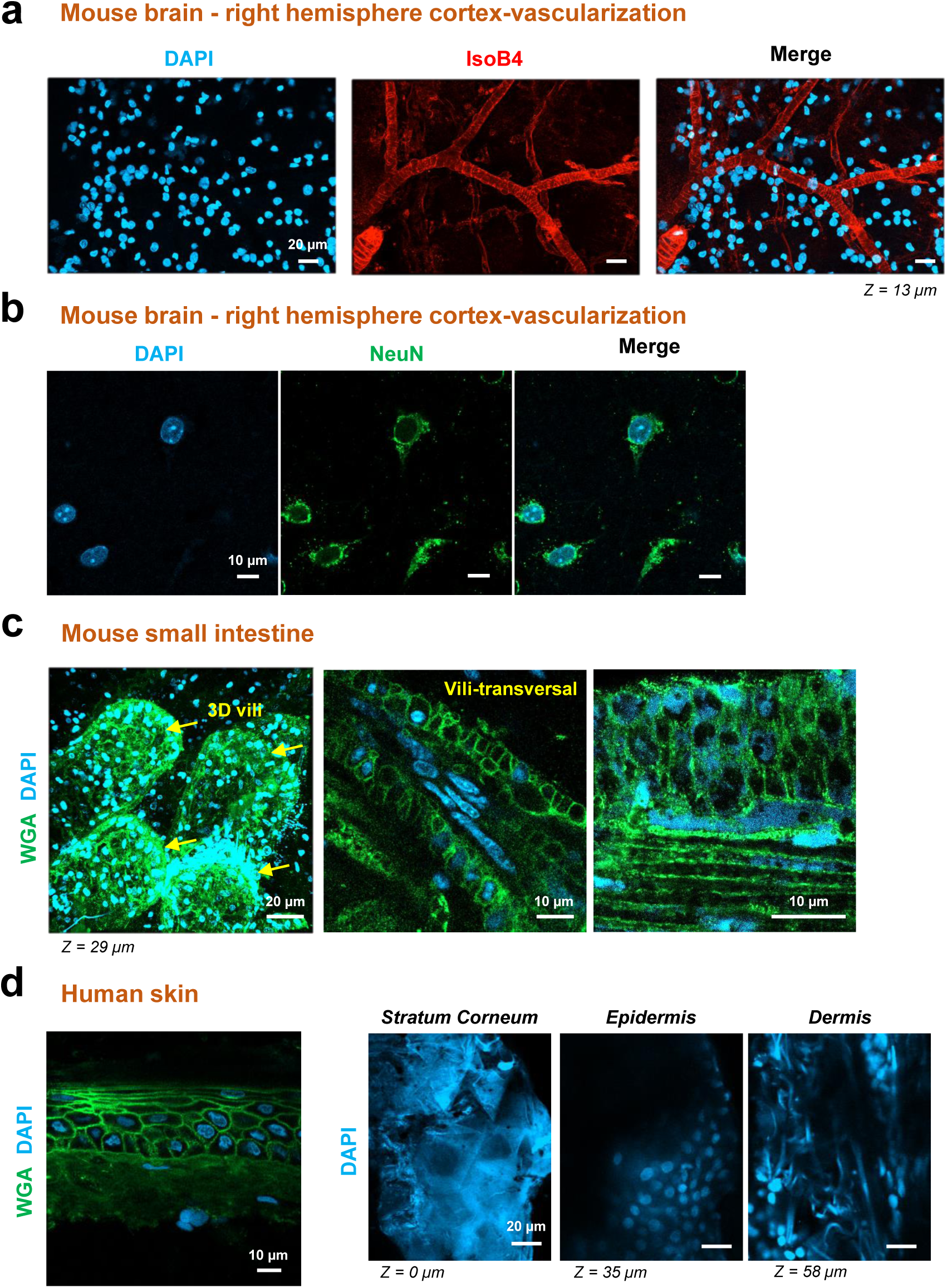
Application of the 3D-NaissI method to various tissues. Representative confocal 2D/3D imaging (2-4 independent experiments) of different tissues following: **a** fluorescent Isolectin-B4 (IsoB4-vascular marker) and DAPI co-staining or **b,** DAPI and NeuN (neuronal marker) in fresh right hemisphere cortex, or **c,** Fluorescent wheat germ agglutinin (WGA) and DAPI co-staining of fresh small intestine from 2-month-old male mice. **d** Fluorescent WGA and DAPI co-staining in fresh human chest skin from a 69-year-old man.

**Supplementary Movie 1. Cardiac Dyadic cleft 3D overall view.** Representative 3D animation of junctophilin-2 (Green) and Ryanodin receptor (Red) expression at the dyadic cleft within cardiomyocytes related to Supplementary Fig. 15b.

